# Alteration in the intracellular Na^+^/K^+^ ratio regulates gene expression through the shift of G-quadruplex dynamics in living cells

**DOI:** 10.1101/2025.10.30.685525

**Authors:** D.A. Fedorov, A.M. Gorbunov, E.G. Maksimov, S.V. Sidorenko, A.M. Varizhuk, A.V. Aralov, O.E. Kvitko, I.Y. Petrushanko, A.A. Makarov, O.D. Lopina, E.A. Klimanova

## Abstract

Alteration in the intracellular Na^+^_i_/K^+^_i_ ratio can play an important regulatory role in many mammalian cells. Here, we demonstrated that the expression of some early response genes does depend on the intracellular Na^+^_i_/K^+^_i_ ratio and that the inhibition of DNA-binding activity of transcription factors c-Fos and c-Jun attenuates the upregulation of *Zfp36* and *Egr1* genes in HeLa cells. Both replacement of intracellular Na^+^ and K^+^ by Li^+^ and the dissipation of Na^+^_i_/K^+^_i_ gradient demonstrated a similar effect on the transcription of *Fos*, *Jun*, *Zfp36*, and *Ptgs2*. Oligonucleotide sequences from the *Fos*, *Kit*, and *Myс* promoters with an ability to fold into G-quadruplexes (G4s) change their conformations differently depending on the ratio of Na^+^, K^+^, and Li^+^ ions *in vitro*. Fluorescence lifetime imaging microscopy studies using DNA G4-specific dye DAOTA-M2 revealed that changes in compositions and ratios of monovalent metal cations in HeLa cells were accompanied by alterations in intranuclear G4s dynamics. The data demonstrate that these structures may be considered as intracellular sensors of monovalent metal cations regulating gene expression through the shift of G4s dynamics.

## Introduction

An imbalance of Na^+^ and K^+^ between the cytoplasm and extracellular medium maintained by Na,K-ATPase (NKA) is the basis for the functioning of any animal cell. The imbalance is essential for the maintenance of the resting membrane potential and the generation of the electrical excitability of cells; it supports secondary active transport of metabolites and ions, and involves in regulation of intracellular pH and cell volume. Normally, the intracellular Na^+^_i_/K^+^_i_ ratio is strictly maintained at a low level. Nevertheless, dissipation of Na^+^_i_/K^+^_i_ gradient can be observed during intensive work of excitable cells and under some pathological conditions. Even short periods of synaptic activity cause a local increase in Na^+^_i_ in apical dendrites from 10 to 30 mM [1]. Intense exercises elevate Na^+^_i_ in human and animal skeletal muscle by 3- to 4-fold, decreasing K^+^_i_ by 15-25% [2,3]. A variety of pathological conditions, including heart failure, ischemia, senescence, Alzheimer’s disease and cancer, can also be accompanied by a significant increase in the Na^+^_i_/K^+^_i_ ratio even in non-excitable cells [4–7]. Thus, intracellular concentrations of monovalent cations, contrary to widespread belief, can undergo serious changes in animal cells.

The imbalance of monovalent cations due to the inhibition of NKA alters gene expression [5]. According to the traditional paradigm, these changes in gene transcription are mediated by a rise in Ca^2+^_i_, which is induced by the increase in the Na^+^_i_ through the operation of Na^+^/Ca^2+^exchanger and/or by operation of voltage-gated Ca^2+^ channels. However, the studies of transcriptomic changes in different cells (HeLa, HUVEC, RVSMC, C2C12) in response to NKA inhibition by ouabain in the presence of extra- and intracellular Ca^2+^ chelators have shown that Ca^2+^ chelators increases rather than decreases the number of genes with altered expression in response to NKA inhibition [8]. The increase in the expression of genes in ouabain-treated cells was also independent on the presence of the inhibitors of voltage-gated Ca^2+^ channels, Na^+^/Ca^2+^ exchanger, calmodulin antagonists, and inhibitors of Ca^2+^/calmodulin-dependent protein kinases and phosphatases [9]. The data indicate the existence of Na^+^_i_/K^+^_i_-sensitive and Ca^2+^-independent mechanism(s) of excitation-transcription coupling [8]. Importantly, many of Na^+^_i_/K^+^_i_-sensitive genes identified by the analysis of transcriptomes of various mammalian cells are transcription factors and translation regulators [8]. It may be assumed that only some of Na^+^_i_/K^+^_i_-sensitive genes are regulated directly by the Na^+^_i_/K^+^_i_ ratio, and the remaining genes have indirect regulation by the upstream Na^+^_i_/K^+^_i_-sensitive factor(s).

Among the previous found Na/K-sensitive genes the *Fos* takes the greatest attention. It encodes a 62 kDa c-Fos protein, which can form heterodimers with c-Jun protein [10], encoded by another Na^+^_i_/K^+^_i_-sensitive gene, *Jun* [11]. The resulting AP1-complex is a transcription factor that regulates gene expression in response to extracellular stimuli [12]. *Fos* is in the "first echelon" of the cellular response that takes place within minutes [13]. Being a classic "immediate early response" gene, which is rapidly activated and rapidly "silenced". The transcriptional regulation of the *Fos* is quite complex, and its promoter includes many regulatory sequences, such as CRE, SRE, SIE, RCE, and TRE, which is the landing site of AP-1 factor, that provides a negative feedback preventing *Fos* overexpression [14].

The lack of Ca^2+^ involvement in *Fos* response gene to the intracellular Na^+^_i_/K^+^_i_ ratio suggests a presence of some molecular sensor(s) of these cations. The search for such a sensor among known protein regulators of the *Fos* promoter has not been successful. Thus, it was shown that the most important regulators of the *Fos* promoter, namely TCF, SRF, CREB, and AP-1, are not sensitive to the intracellular Na^+^_i_/K^+^_i_ ratio [15]. To localize the regions of the *Fos* promoter that are involved in gene regulation by monovalent cations, Nakagawa et al. used genetic constructs containing the human *Fos* under the control of 5’-truncated promoters with deletions [16]. As a result, Na^+^_i_/K^+^_i_ sensitivity of the *Fos* promoter was conferred to the participation of the SRE element, and to the region from −222 to −123 b.p. relative to the transcription start point.

We hypothesize that the promoters of Na^+^_i_/K^+^_i_-sensitive genes can perceive intracellular concentrations of monovalent cations through the folding/unfolding of G-quadruplexes (G4s) that are noncanonical secondary structures of nucleic acids. The monomeric unit of G4 is a G-tetrad, 4 guanine bases linked by Hoogsteen-type hydrogen bonds. G4-structures are topologically very diverse: depending on the orientation of nucleic acid chains, G4s can take parallel, antiparallel, or hybrid topological forms [17]. The most important contribution to the stabilization of G4 provides with the specific binding of metal cations [18,19]. The data on the melting of G4s give a following ranks of cation efficiency of G4s stabilization: K^+^ > Rb^+^ > Na^+^ > Li^+^ = Cs^+^ [20]. Thus, the predominant K^+^ cations in living cells generally support G4s formation, which explains the widespread occurrence of G4s in animal and human genomes [21].

Bioinformatic algorithms detect more than 700,000 consensus sequences in the human genome that can form G4s [22]. G4s are found in 42.7% of human gene promoters, and their occurrence is particularly high near the transcription start region [23]. These structures can be involved in creating binding sites for a number of different proteins. In addition, G4s can prematurely terminate transcription in the case of intermolecular G4 formation between the DNA coding strand and the RNA transcript during its synthesis by RNA polymerase [24]. Bioinformatic analysis of the *Fos* 5’-flanking region sequence from −550 b.p. to +155 b.p. relative to the transcription start point predicts the formation of at least 3 G4s within it, one of them is a part of a region sensitive to the intracellular Na^+^_i_/K^+^_i_ ratio found by Nakagawa et al. [16]. This may be in favor of the involvement of these G4s in the regulation of *Fos* by monovalent metal cations.

Herein we show that *Fos* is a regulator that restrains the overexpression of Na^+^_i_/K^+^_i_-dependent genes triggered by monovalent cations perturbations. We find that the tested oligonucleotide sequences within the *Fos* promoter form G4s DNA *in vitro* and have different sensitivities to monovalent cations. Finally, we demonstrate that monovalent cations imbalance affects the dynamics of G4s DNA in the nuclei of HeLa cells. These data allow us to suggest that intracellular sensors of monovalent metal cations are G4s.

## Methods

### Cell culturing

HeLa cells were maintained in Dulbecco’s Modified Eagle Medium (DMEM, PanEco, Russia) supplemented with 10 % fetal bovine serum (FBS, Cytiva, USA) and 100 U/ml penicillin and 100 mg/ml streptomycin (Gibco, USA) in a humidified atmosphere with 5 % CO_2_/balance air at 37 °C. The growth medium was changed every three days. After reaching 80-90% confluency, cells were seeded onto 12-well plates (100000 cells per well) and cultured up to 80-90% confluency. To establish quiescence, cells were incubated for 24 hours in DMEM supplemented with 0.1% FBS, after which the cells were subjected to experimental treatments. To invert the intracellular [Na^+^]_i_/[K^+^]_i_ ratio, cells were incubated in the presence of 3 μM ouabain (Sigma Aldrich, USA) for 3 h. To stabilize intracellular G4s, cells were incubated in the presence of 10 μM pyridostatin (Cayman Chemical, USA) for 24 h. To inhibit the DNA-binding activity of FOS, cells were incubated in the presence of 20 μM T-5224 (ApexBio Technology, USA) for 24 h.

In the study of Li^+^ effect on gene expression, cells were incubated in the presence of Li-medium (10 mM HEPES-LiOH (pH 7.4), 130 mM LiCl, 5.2 mM KCl, 1 mM CaCl_2_, 0.5 mM MgCl_2_, 0.4 mM MgSO_4_, 0.3 mM KH_2_PO_4_, 6 mM glucose) for 4 h. As a control in these experiments, we used Na-medium, which is the same composition as Li-medium with all Li^+^ ions replaced by Na^+^.

### Intracellular ions content measurement

12-well plates were transferred onto ice, experimental medium was quickly removed and HeLa cells were washed three times with 3 ml of an ice-cold 0.1 M MgCl_2_ solution in doubly deionized (DDI) water. Then, 1.5 ml of 5% trichloroacetic acid (TCA) in DDI water was added to each well, followed by incubation at 4 °C overnight for complete extraction of ions from cells. Cell precipitates were suspended and centrifuged during 5 min at 15000 g. Supernatants were transferred into test tubes and stored at −20°C. Then cell precipitates were resuspended in 0.75 ml of 0.1 M NaOH and incubated at 65 °C during 1 h for complete protein dissolving. Resulted protein solutions were used for protein amount quantification by Lowry protein assay [25]. The Na^+^_i_, K^+^_i_, and Li^+^_i_ contents in TCA extracts were measured by flame atomic absorption spectrometry using the Kvant-2m1 spectrometer (Cortec, Russia) with propane-air mixture at 589 nm, 766.5 nm, and 670.8 nm, respectively. KCl (0.5–4 mg/L K^+^), NaCl (0.05–2 mg/L Na^+^), and LiCl (0.25–4 mg/L Li^+^) solutions in 5% TCA in DDI water were used for calibration. The Na^+^_i_, K^+^_i_, and Li^+^_i_ contents in each well were normalized on protein amount in the same well.

### Cell viability measurement

The AlamarBlue assay [26] was carried out according to the manufacturer’s instructions. Briefly, experimental medium was removed and wells were filled with 1.5 ml of fresh experimental medium. 45 μl of AlamarBlue reagent were added to this volume of medium in each well and mixed gently. Following 2 h of incubation fluorescence of medium was measured at the excitation and emission wavelength of 570 and 585 nm, respectively, using a Synergy H4 Multi-Mode Hybrid Microplate Reader (BioTek, USA). For each experiment, wells containing only medium with AlamarBlue reagent without cells were also prepared and incubated during 2 h for quantification of background fluorescence. Fluorescence signals from each well were normalized on protein amount in the same well, which was determined by Lowry protein assay [25]. Viability of control wells was referred to as 100 %.

### HeLa cells nuclei isolation

Isolation of HeLa cells’ nuclei was performed as described in [27]. HeLa cells were cultured in a culture flask (Corning, USA) with a cell growth surface of 75 cm^2^ up to 90 % confluence, after which the medium was discarded, and the cells were washed with 15 ml of a Hanks’ solution without Ca^2+^, Mg^2+^ ions and phenol red (137 mM NaCl, 5.4 mM KCl, 0.42 mM Na_2_HPO_4_, 0.44 mM KH_2_PO_4_, 5.6 mM glucose) and then 4 ml of hypotonic buffer was added (10 mM HEPES-NaOH pH 7.9, 1.5 mM MgCl2, 1 mM KCl, 0.5 mM DTT). After that, the cells were detached using a special scraper and transferred to a Downs homogenizer (glass-glass). Homogenization was carried out on ice by 30 rotational and translational movements of the pestle. The resulting suspension of destroyed cells was distributed into Eppendorf-type tubes and centrifuged at 700 g and +4 °C for 5 minutes. The supernatant was discarded, and the precipitate containing HeLa nuclei was used for experimental purposes.

### Intranuclear ions content measurement

HeLa nuclei were isolated in Eppendorf-type tubes as described earlier. The mass of precipitate containing nuclei (m) was measured using Ohaus PA214C analytical balance (Ohaus, USA). The nuclei were then put in one of the four incubation media listed in Table 1 by adding a comparable volume of one or another medium to each precipitate (V_x_). The sediments were suspended in these 4 media and the final volume of each sample was determined by pipetting (V_y_). The mean density of isolated nuclei was determined using the following simple equation: 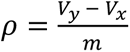, averaging the result for 4 samples. Each of four nuclei suspensions then was distributed into 6 separate tubes, and the volume of the incubation medium in each sample was adjusted to 1 ml. The samples were incubated at 37 °C for 15 min, after which the non-nuclear medium was disposed by centrifugation at 8000 g and 4 °C for 2 min. The tubes were then centrifuged under the same conditions again and the remnants of the supernatant were removed using a pipette with a retracted tip. Next, the mass of the nuclear sediment in each sample (m_i_) was determined using Ohaus PA214C analytical balance and 1.5 ml of 5 % aqueous TCA were added in order to destroy the nuclei. The content of Na^+^ and K^+^ in the obtained nuclear lysate was determined with atomic absorption spectrometry similar to experiments with HeLa cells. The volume of the nuclear fraction in each sample was calculated as 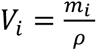, using the mean density of HeLa nuclei in a precipitate (p). Thus, the concentrations of Na^+^ and K^+^ ions inside isolated HeLa nuclei was calculated by normalizing the content of these ions by the volume of the nuclear fraction.

**Table 1.** Composition of media used for incubation of isolated HeLa nuclei.

| Media type | Buffer | NaCl и KCl | Ouabain |
| --- | --- | --- | --- |
| I | 20 mM HEPES-imidazole (pH 7,8)<br>5 mM MgCl<sub>2</sub><br>0,5 mM DTT<br>0,25 M sucrose<br>1 mM EGTA | 100 mM KCl,<br>10 mM NaCl | — |
| II |  |  | 1 μM |
| III |  | 10 mM KCl,<br>100 mM NaCl | — |
| IV |  |  | 1 μM |

Similar experiments were carried out using HeLa nuclei suspended into two different volumes (150 µl and 1.5 ml) of incubation media of type I and III from Table 1.

### Real-time quantitative RT-PCR

HeLa cells (∼1×10^6^ cells) were washed with an ice-cold Mg^2+^/Ca^2+^-free HBSS buffer (137 mM NaCl, 5.4 mM KCl, 0.42 mM Na_2_HPO_4_, 5.5 mM glucose, 4.16 mM NaHCO_3_) and 400 *μ*l of Trizol reagent was added in order to isolate the total RNA. After isolation of the aqueous phase containing nucleic acids using chloroform and treatment with 96 % ethanol, further steps of RNA isolation and treatment with DNase were performed on Quick-RNA MicroPrep columns. The ImProm-II^TM^ Reverse Transcription System kit was used for the reverse transcription reaction. In all cases, procedures were followed according to the manufacturer’s instructions. The real-time PCR was carried out with the Bio-Rad Real-Time PCR System (Bio-Rad, California, Hercules, CA, USA). Primers (Syntol, Moscow, Russia, Table 2) were added to a final concentration of 200 nM. Amplification modes: 95 °C 5 min; 95 °C 10 s; 58 °C 17 s; 72 °C 20 s; 40 repetitions of steps 95 °C 10 s, 58 °C 17 s, 72 °C 20 s; melting curve from 72 to 95 °C, increment 0.5 °C 5 s. The selection of primers was performed using the NCBI and BLAST search databases. The expression level of each gene of interest was calculated using the reference *Rplp0* (Ribosomal Protein Lateral Stalk Subunit P0) gene by the 2^-ΔΔСt^ method [28]. Gene expression in the control was taken as 100 %. To verify the PCR products, they were sequenced. Electrophoresis was performed in 2 % agarose gel using TBE buffer in the presence of ethidium bromide for ∼1h at 75 V. PCR products were extracted from the gel using a QIAquick Gel Extraction Kit according to the manufacturer’s protocol. DNA sequencing was carried out by Genom (Moscow, Russia). The sequences were aligned using the BLAST program. All DNA sequences were as stated.

**Table 2.**
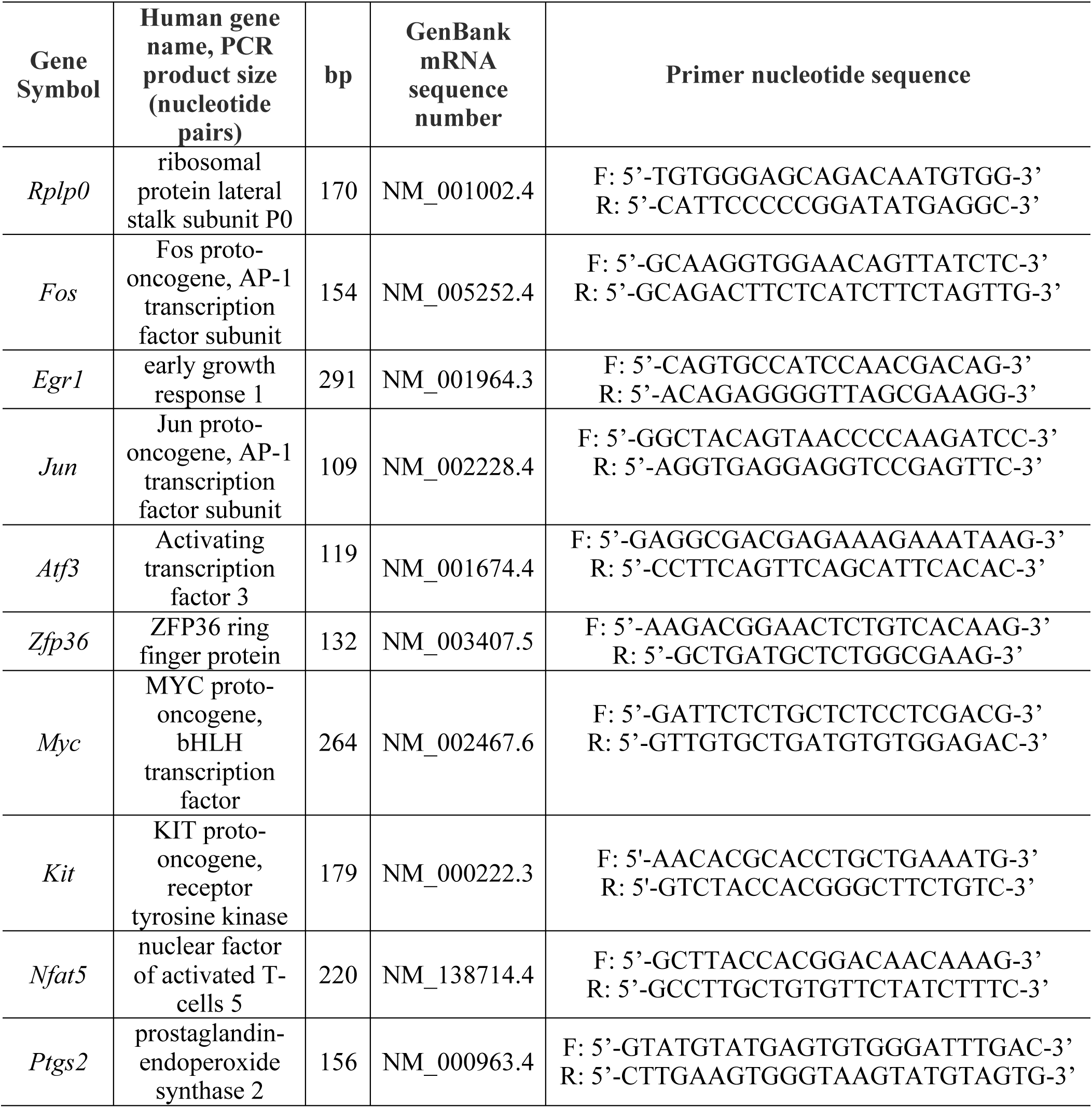
Primer sequences used in this study.

### Circular dichroism

Potential G-quadruplex-forming sequences (PQS) of the human *Fos* promoter (FOS_G4_1, FOS_G4_2, FOS_G4_3) were predicted by G4catchall algorithm [29]. A modified G4-forming sequence of MYC promoter [30], a human telomeric G-quadruplex 22AG_G4_ and a G-quadruplex of KIT promoter [31] were used as control oligonucleotides that are known to form G-quadruplex *in vitro* (Table 3). Circular dichroism spectra of preannealed oligonucleotide solutions (2 μM) in 10 mM HEPES-Tris buffer, pH 7.4, supplemented with NaCl, KCl, and/or LiCl to a final total concentration of Na^+^, K^+^, and Li^+^ of 100 mM were registered using a Chirascan spectrophotometer (Applied Photophysics, UK) and standard quartz cuvettes of 1 cm path at room temperature at wavelength 230 – 330 nm. For each oligonucleotide, three measurements were averaged after background subtraction.

**Table 3.**
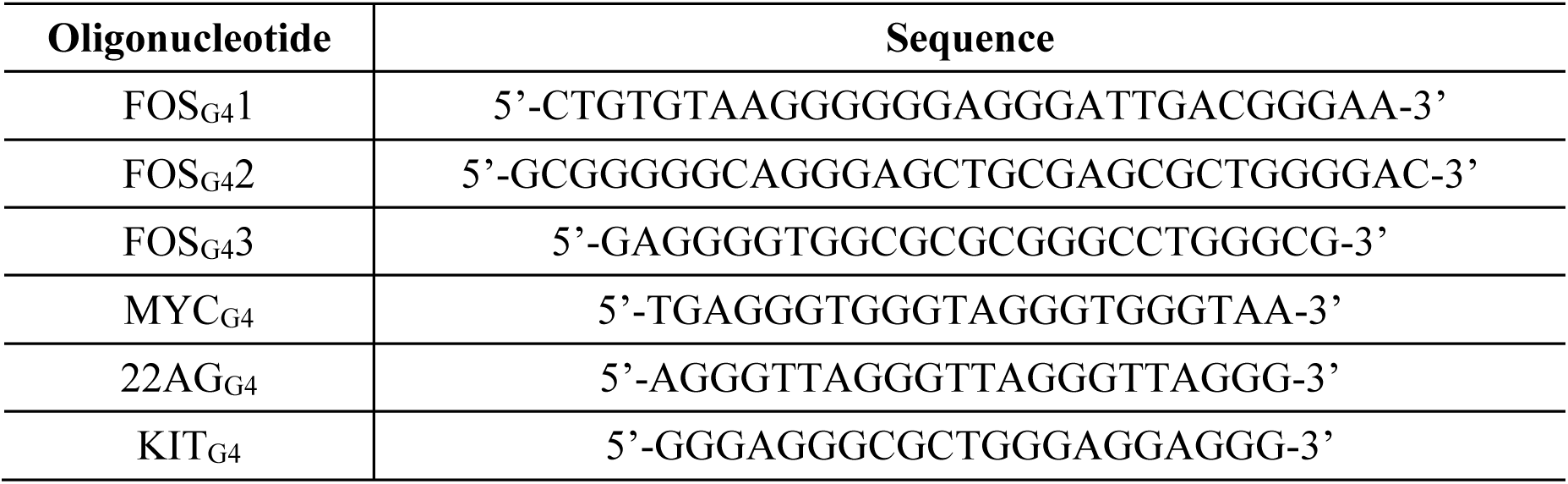
Oligonucleotides used in circular dichroism analysis.

### In vitro and in cellulo DAOTA-M2 fluorescence lifetime measurements

DAOTA-M2 was synthesized as previously reported [32], ^1^H NMR and MS data are consistent with published data (Fig. S1, S2). *In vitro* evaluation of DAOTA-M2 fluorescence lifetime was conducted using fluorescence lifetime imaging microscopy (FLIM). 1 μM oligonucleotides (Table 3) were annealed in 10 mM HEPES-Tris pH 7.4 supplemented with NaCl, KCl or LiCl as mentioned previously. Then, 1 μM DAOTA-M2 was added to the samples. FLIM was performed in a 96-well plate.

For *in cellulo* experiments, HeLa cells were incubated in DMEM supplemented with 0.1 % FBS and 20 μM DAOTA-M2 for 24 h. Before microscopy, the medium was changed to HBSS (untreated cells and cells treated by ouabain) or to Na-/Li-medium with 20 μM DAOTA-M2 for 30 min or 4 h, respectively.

FLIM was conducted at room temperature (24 °C) and in atmospheric air. Fluorescence lifetime images were obtained using the time-correlated single-photon-counting system SPC150 and the confocal galvano scanner DCS-120 (Becker&Hickl, Berlin, Germany) installed on the Eclipse Ti2 (Nikon, Japan) microscope as described in [33]. Excitation was performed with a 473 nm picosecond laser BDS-SM-473-LS-101 (Becker&Hickl) at 20 MHz. Detection was performed using a 590 nm bandpass filter with 40 nm band width (Chroma, Bellows Falls, VT, USA). For *in cellulo* experiments, a A Plan-Apochromat 60× 1.4 NA lens (Nikon, Japan) was used to collect images at 512 × 512 pixel resolution for 1000 s, for *in vitro* experiments, a 20× 0.75 NA lens (Nikon, Japan) was used to collect fluorescence of solutions in a 96-well plate, in such case scanning mode was used to prevent bleaching at single point, but adjusted to 16 × 16 pixel resolution and integration for 120 s. FLIM data was processed via SPCImage (Becker&Hickl) as described in [33]. Traces were fitted to a bi-exponential decay function I(t) = *I*_0_(*a*_1_*e*^-t/t1^ + *a*_2_*e*^-*t*/*t*2^) where *a_1_* and *a_2_* are variables normalized to unity, and the intensity-weighted average lifetime (τ_w_) was calculated using the equation:

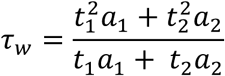

### Statistics

Statistical data analysis was performed using the GraphPad Prism version 10.4.0 software. All sample distributions were checked for normality using the Shapiro-Wilk test. To compare two independent groups, the Student’s t-test was used for data with homogeneous variances as well as the Welch’s t-test when the assumption of homogeneity of variance was not met. Multiple comparisons were performed using single-factor analysis of variance (ANOVA) followed by the Tukey’s test (for data with homogeneous variances) or Welch-corrected ANOVA followed by post hoc Dunnett’s T3 test (when the assumption of homogeneity of variance is not met). Homogeneity of variance in all cases was checked with Brown-Forsythe and Bartlett’s test. The details of statistical analysis are given in the figure legends.

## Results

### c-Fos/c-Jun attenuates Na^+^_i_/K^+^_i_-dependent expression of Zfp36 and Egr1 genes at mRNA level

The targets of our study were the following genes: *Fos*, *Jun*, *Zfp36*, *Egr1*, *Atf3*, *Ptgs2*, *Kit*, *Myc*, and *Nfat5*. Among them, *first six* are Na^+^_i_/K^+^_i_-sensitive genes [8]; *Kit* and *Myc* are genes whose transcription is regulated involving G-quadruplexes located in their promoters [24]; *Nfat5* encodes an osmoprotective transcription factor that is insensitive to intracellular Na^+^_i_/K^+^_i_ ratio [34].

Since previously we had found using bioinformatic approaches, that c-Fos can regulate some of Na^+^_i_/K^+^_i_ -sensitive genes [35], we at first studied the effect of T-5224, an inhibitor of the DNA-binding activity of c-Fos/c-Jun complex [36], on the expression of the tested genes. To exclude the effect of monovalent cations imbalance on gene expression in HeLa cells pre-incubated with T-5224, we tested its effect on ion transport in these cells. As shown in Figure 1a, treatment of HeLa cells with 20 µM T-5224 for 24 h had no significant effect on the intracellular Na^+^_i_ and K^+^_i_ content. Incubation of the cells with 3 µM ouabain for 3 h provided an increase of intracellular Na^+^_i_ (from 73 ± 4 to 879 ± 127 nmol/ mg protein) and a decrease of intracellular K^+^_i_ (from 1015 ± 62 to 223 ± 45 nmol/ mg protein). We also did not detect any effect of T-5224 on the content of these cations in ouabain treated cells (Fig. 1a).

**Fig. 1.**
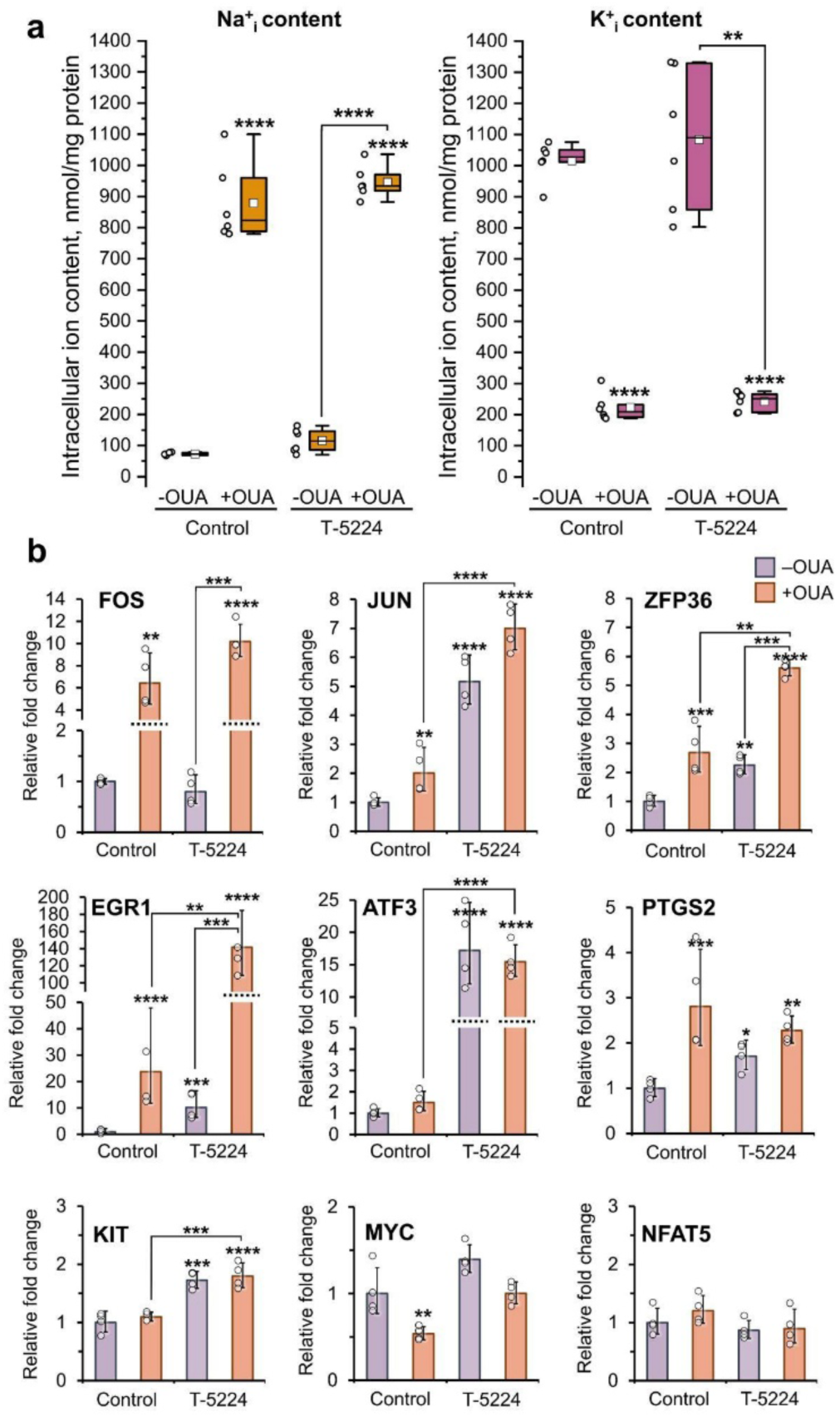
Na^+^_i_/K^+^_i_-dependent gene upregulation in HeLa cells does not rely on c-Fos/c-Jun DNA-binding activity. **a.** Box plot diagrams showing the intracellular Na^+^ and K^+^ content in HeLa cells subjected to 3 h inhibition of Na,K-ATPase with 3 µM ouabain or in ouabain-free conditions after 24 h incubation in control (DMEM) or in the presence of 20 µM T-5224, an inhibitor of the DNA-binding activity of c-Fos/c-Jun. Data are from 6 biological replicates (n=6), line represents median, square represents mean, box represents 25–75th percentile, whiskers represent range within 1.5 IQR. **b.** Expression levels of genes in HeLa cells under conditions described in **a**. Bar chart represents geometric mean relative fold change and standard deviation (SD). Data are provided from 4 biological replicates (n=4). The reference gene was *Rplp0*. Fold changes are relative to ouabain-free control. Statistically significant differences were revealed using Welch-corrected ANOVA followed by post hoc Dunnett’s T3 test (for data in **a**) or ANOVA followed by post hoc Tukey’s test (for data in **b**); \**p < 0.05,* \*\**p < 0.01*, \*\*\**p < 0.001*, \*\*\*\**p < 0.0001*.

Then, we examined the effect of c-Fos/c-Jun inhibitor on the transcription of all above mentioned genes. As shown in Figure 1b, in the case of *Jun, Zfp36, Egr1*, *Atf3*, *Ptgs2*, and *Kit* genes, similar expression pattern was observed: incubation of cells in the presence of 20 µM T-5224 for 24 h resulted in the increase of the amount of mRNA encoded by these genes in comparison with the control samples (5-, 2-, 10-, 15-, 1.5-, and 2-fold, respectively), whereas the amount of *Fos, Myc,* and *Nfat5* mRNA was unchanged. Treatment of cells with 3 µM ouabain increased the amount of mRNA of *Fos*, *Jun*, *Zfp36*, *Egr1*, and *Ptgs2* (6-, 2-, 2.5-, 20-, and 3-fold, respectively), whereas 2-fold decreased the amount of *Myc* mRNA. Under these conditions, the expression of *Atf3*, *Kit,* and *Nfat5* remained unchanged compared to control samples. Ouabain also increased the amount of mRNA of *Jun*, *Zfp36*, *Egr1*, *Atf3*, *Kit* (3.5-, 2-, 7-, 15-, and 2-fold, respectively) and had no effect on the expression of *Fos*, *Ptgs2, Myc,* and *Nfat5* in T-5224-treated cells in comparison with cells incubated with ouabain alone. The data allow us to draw the following conclusions. <u>First</u>, c-Fos/c-Jun complex limits the mRNA levels of *Jun*, *Zfp36*, *Egr1*, *Atf3*, *Ptgs2*, and *Kit* genes in untreated cells. <u>Second</u>, c-Fos/c-Jun is not involved in Na^+^_i_/K^+^_i_-dependent gene upregulation of *Fos*, *Zfp36*, and *Egr1* in HeLa cells. <u>Third</u>, c-Fos/c-Jun attenuates Na^+^_i_/K^+^_i_-dependent regulation of *Zfp36* and *Egr1* mRNA levels.

### Expression of Fos, Jun, Zfp36, and Ptgs2 genes is affected by Li^+^ ions but not by pyridostatin

We hypothesize that DNA G4s may be involved in the control of Na^+^_i_/K^+^_i_-dependent gene expression. It has been shown that Li^+^ ions have a destabilizing effect on conformation of DNA G4s *in vitro* [21]; therefore, we decided to evaluate the effect of these cations on the expression of Na^+^_i_/K^+^_i_-dependent genes. In order to load cells with Li^+^, HeLa cells were incubated for 4 h in the Li-medium (Fig. 2a), in which Na^+^ ions were replaced with Li^+^ ones [37,38]. Cells incubated in a similar media containing Na^+^ ions (Na-medium) were used as control samples (Fig. 2a). Incubation of cells in Li-medium decreased both intracellular Na^+^_i_ from 97 ± 28 to 8 ± 2 nmol/mg protein and intracellular K^+^_i_ from 1127 ± 111 to 258 ± 82 nmol/mg protein (Fig. 2b). Intracellular Li^+^_i_ under these conditions amounted to 546 ± 64 nmol/mg protein (Fig. 2b). As shown in Figure 2c, the replacement of extracellular Na^+^ with Li^+^ ions resulted in an increase of mRNAs level of *Fos*, *Jun*, *Zfp36,* and *Ptgs2* genes (9-, 10-, 4-, and 1.5-fold, respectively) and a decrease in *Kit, Myc,* and *Nfat5* mRNAs level (1.25-, 17-, and 1.5-fold, respectively), but did not affect *Egr1* and *Atf3* expression.

**Fig. 2.**
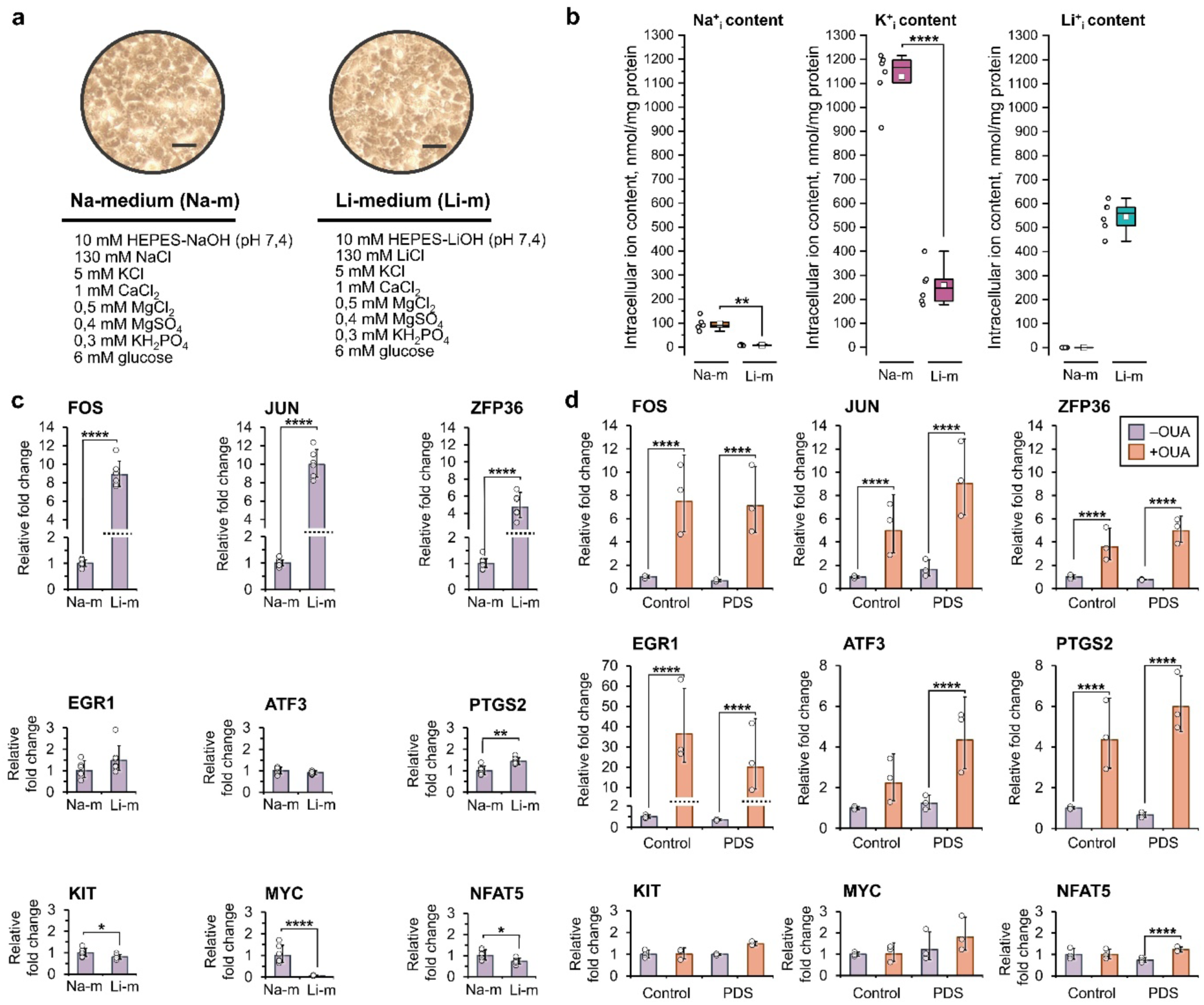
Li^+^ ions but not pyridostatin upregulates the expression of some Na^+^_i_/K^+^_i_-sensitive genes. **a.** Representative images of HeLa cells incubated in Na- or Li-medium for 4h. Scale bars are 5 µm. **b.** Boxplot diagrams showing the intracellular Na^+^, K^+^ and Li^+^ content in HeLa cells incubated in Na- or Li-medium for 4h. Data are from 5-6 biological replicates (n= 5-6), line represents median, square represents mean, box represents 25–75th percentile, whiskers represent range within 1.5 IQR. **c.** Effect of 4 h incubation in Li-medium on gene expression in HeLa cells compared to Na-medium. Bar chart represents geometric mean relative fold change and standard deviation (SD). Data are provided from 6 biological replicates (n=6). The reference gene was *Rplp0*. Fold changes are relative to Na-medium. **d.** Expression levels of genes in HeLa cells subjected to 3 h inhibition of Na,K-ATPase with 3 µM ouabain or in ouabain-free conditions after 24 h incubation in control (DMEM) or in the presence of 10 µM pyridostatin (PDS). Bar chart represents geometric mean relative fold change and standard deviation (SD). Data are provided from 3 biological replicates (n=3). The reference gene was *Rplp0*. Fold changes are relative to ouabain-free control. Statistically significant differences were revealed using Welch-corrected or Student’s t-test (for data in **b** and **c**) or ANOVA followed by post hoc Tukey’s test (for data in **d**); \**p < 0.05,* \*\**p < 0.01*, \*\*\**p < 0.001*, \*\*\*\**p < 0.0001*.

Taking into account the fact that pyridostatin stabilizes G4s [39], we evaluated the action of pyridostatin on the mRNA level of tested genes under conditions of the monovalent metal cations gradient dissipation. We found no effect of 10 µM pyridostatin on the monovalent metal cations content in HeLa cells (Table S1), whereas incubation of the cells in the presence of 3 µM ouabain for 3 h led to the increase in the intracellular Na^+^_i_/K^+^_i_ ratio from ∼0.1 to ∼4 compared to control samples (Fig. 1a). Figure 2d shows that pyridostatin did not affect the mRNA levels of the tested genes in either untreated or ouabain-treated cells, whereas the effect of ouabain is retained in pyridostatin-treated cells.

Thus, the expression of *Fos*, *Jun*, *Zfp36,* and *Ptgs2* genes depends on both the changes in Na^+^_i_/K^+^_i_ ratio and the presence of Li^+^ ions inside the cell, i.e. it demonstrates monovalent metal cation-dependent regulation.

### DNA G4s demonstrates different sensitivity to the ratio of monovalent metal cations in vitro

Among the genes showing monovalent metal cation-sensitive expression, *Fos* deserves special attention. First, we did not detect c-Fos/c-Jun-dependent negative feedback regulation of *Fos* expression under conditions of dissipation of the monovalent cation gradient (Fig. 1b). Second, it was previously shown that the amount of *Fos* mRNA correlates with the value of the intracellular Na^+^_i_/K^+^_i_ ratio [11,15]. Third, changes in *Fos* expression are observed already in the initial steps of the dissipation of Na^+^ and K^+^ ion gradient [11,15]. We therefore suggest that G4s within the *Fos* promoter may provide such regulation. Probably, this mechanism of regulation is not limited to *Fos* and may extend to other genes.

G4s are a promising candidate for the role of monovalent cation sensors and switch. A curious fact is their extremely high frequency in the promoters of proto-oncogenes. The *Myc* and *Kit* genes have long been considered as classic models of genes containing G4s in their promoters [24]. Although *Fos* is also classified as a proto-oncogene, the data concerning the presence of G4s in the promoter of this gene were published relatively recently [40]. We also conducted a bioinformatics analysis of the human *Fos* promoter region and identified at least three potential G-quadruplex-forming sequences (PQSs) (FOS_G4_1. FOS_G4_2, FOS_G4_3).

Next, we synthesized oligonucleotides corresponding to the PQS in the *Fos* gene promoter (Fig. 3a) and evaluated using circular dichroism (CD) spectroscopy the structure of these oligonucleotides in the presence of following K^+^/Na^+^ ratios: 100 mM : 0 mM, 80 mM : 20 mM, 50 mM : 50 mM, 20 mM : 80 mM, and 0 mM : 100 mM. In addition, we used a medium containing 2 mM Na^+^, 40 mM K^+^, and 80 mM Li^+^. This composition reflects the concentrations of monovalent cations in HeLa cells when extracellular Na^+^ ions are partially replaced by Li^+^ ions. Besides the *Fos* gene, we used G4 from the *Kit* and *Myc* promoters, as well as 22AG, which is the G4 of human telomeric DNA [24,31].

**Fig. 3.**
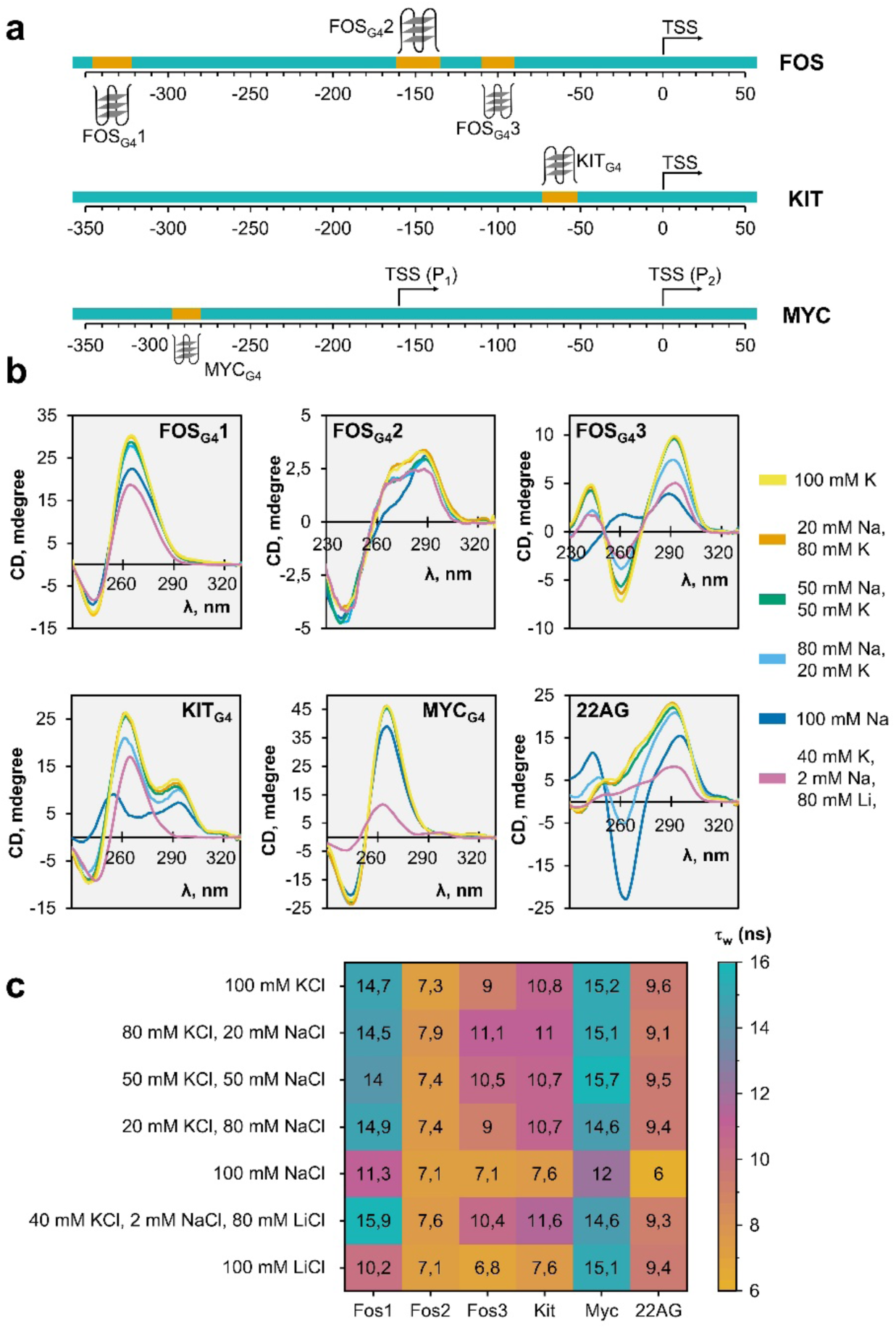
Promoter G4s structure are sensitive to monovalent metal cation perturbations *in vitro*. **a.** Schematic image of the *Fos*, *Kit*, and *Myc* promoters. DNA sites forming G4 sequences are marked with color and symbols, located above (for + strand G4) or below (for – strand G4) the line. **b.** Circular dichroism spectra of oligonucleotides forming G-quadruplexes from the *Fos*, *Kit*, and *Myc* promoters and telomeric G-quadruplex (22AG). The oligonucleotides were annealed in presence of 0-100 mM KCl, 0-100 mM NaCl and 0-80 mM LiCl. **с.** A heatmap of τ_w_ of DAOTA-M2 mixed with oligonucleotides forming G-quadruplexes from the *Fos*, *Kit*, and *Myc* promoters as long as telomeric G-quadruplex (22AG) in the presence of 100-0 mM KCl, 0-100 mM NaCl and 0-100 mM LiCl.

First, it should be noted that all the sequences which we studied had spectra characteristic of formed G4s in the presence of 100 mM K^+^, but these G4s formed different types of structures (Fig. 3b). <u>FOS_G4_1</u> has a a parallel G4 structure in all media used. Increasing the Na^+^/K^+^ ratio in the solution slightly affected the CD spectra, whereas replacing K^+^ with Na^+^ resulted in a decrease in peak amplitude by approximately 30%; the presence of Li^+^ reduced peak amplitude by 40%.

The spectrum of <u>FOS_G4_2</u> corresponded to a hybrid G4 in all solutions. A significant change in the spectrum was observed only in the presence of 100 mM Na^+^ and manifested itself as a decrease in ellipticity at 260 nm to 0 millidegrees. The spectrum of <u>FOS_G4_3</u> in the presence of K^+^ ions corresponded to an antiparallel G4 structure. Increasing the Na^+^/K^+^ ratio to 80 mM:20 mM resulted in a 20-50% decrease in peak amplitude; the presence of Li^+^ in the medium had a similar effect. In the presence of 100 mM Na^+^ alone, the spectrum was not characteristic of G4 structures.

<u>KIT_G4_</u> formed a hybrid G4 in the presence of K^+^ and in the absence of Li^+^. The peak amplitude in its spectrum decreased by 20% with increasing the Na^+^/K^+^ ratio to 80 mM:20 mM. In the presence of Na^+^ alone, the spectrum acquired properties uncharacteristic of G4. In the presence of Li^+^, G4 had a parallel structure.

The spectrum of <u>MYC_G4_</u> corresponded to a parallel G4. Complete replacement of K^+^ with Na^+^ reduced the peak amplitude by 15%, and the presence of Li^+^ reduced it by 80%. At a Na^+^/K^+^ ratio of 50 mM:50 mM or less, the spectrum of <u>22AG</u> corresponded to a hybrid G4 structure. In the presence of Li^+^, the peak amplitude dropped by 75%. Increasing the Na^+^/K^+^ ratio to 80 mM:20 mM and replacing K^+^ with Na^+^ led to a change in the G4 type to antiparallel.

Thus, all the oligonucleotides studied, including three PQSs from the *Fos* promoter, formed different types of G4 structures *in vitro*. All these sequences are characterized by different sensitivities to the concentration ratios of monovalent cations.

To evaluate the effect of the different monovalent cations on the of G4s in living cells, we used the fluorescent dye DAOTA-M2 which has an averaged *K_d_* of 1.0 and 1.7 µM for G4s and double-stranded DNA, respectively [41], but has higher fluorescence lifetimes when bound to G4s DNA structures (8-12 ns) compared to double stranded DNA (5-7 ns) [42]. Recently, DAOTA-M2 has successfully been used to visualize G4s DNA dynamics by FLIM in living cells [43].

Since the cation ratio can affect not only the number of G4s but also their topology (Fig. 3b), we decided at first evaluate the effect of different concentrations of monovalent cations on the intensity-weighted average lifetime (τ_w_) of DAOTA-M2 bound to different G4s *in vitro* (Fig. 3c). Thus, we observed distributions of τ_w_ values and determined their maximum at different Na^+^/K^+^ ratio in the case of parallel G4s FOS_G4_1 (11,2-14,9 ns) and MYC_G4_ (12-15,7ns). These values decreased in the following range: KIT_G4_ (7,6-11 ns), FOS_G4_3 (7-11,1 ns), FOS_G4_2 (7-7,9 ns), and 22AG (6-9,6 ns). Importantly, no correlation was found between τ_w_ of DAOTA-M2 bound to the investigated G4s value and the value of the Na^+^/K^+^ ratio, with the only exception being FOS_G4_3. In this case, the parameter had larger values when the Na^+^/K^+^ ratio was low (Fig. 3c). Interestingly, KIT_G4_, FOS_G4_2, and 22AG are characterized by a hybrid topology, while FOS_G4_3 was antiparallel. However, replacing K^+^ with Na^+^ is accompanied with changes in the CD spectral characteristic of all studied G4s *in vitro* and a decrease in τ_w_ (DAOTA-M2). To conclude *in vitro* section, τ_w_ of DAOTA-M2 might depend not only on the number of G4s, but also on their topology.

If we compare the effects of 100 mM K^+^ and 100 mM Na^+^, the investigated G4s τ_w_ (DAOTA-M2) are characterized by a lower value in the presence of Na^+^ ions. Interestingly, no such pattern was found when comparing 100 mM K^+^ and 100 mM Li^+^ (Fig. 3c). However, τ_w_ upon binding to different G4s in the solution containing 40 mM K^+^, 2 mM Na^+^, 80 mM Li^+^ changes differently compared to the data obtained in the solution with 80 mM K^+^ and 20 mM Na^+^: thus, in the case of 22AG, KIT_G4_, FOS_G4_1 this parameter increases, whereas in the case of MYC_G4_, FOS_G4_2, FOS_G4_3 the lifetime of DAOTA-M2 fluorescence decreases. According to CD data (Fig. 3b), in the presence of 40 mM K^+^, 2 mM Na^+^, 80 mM Li^+^, KIT_G4_ structure altered from the hybrid to the parallel type, which is also reflected in the change of τ_w_ (DAOTA-M2) from 111 to 11,6 ns. Thus, these observations may further support the fact that DAOTA-M2 has a greater selectivity to the parallel G4s.

### Monovalent metal cations perturbations influence DNA G4s dynamics in HeLa nuclei

Since we observed that monovalent metal cations perturbations affect both gene expression *in cellulo* and the structure of DNA G4s *in vitro*, we can suggest that Na^+^_i_/K^+^_i_-dependent regulation of gene expression in mammalian cells may be mediated by the changes in the DNA G4s dynamics. The question is risen, how do cytoplasmic and intranuclear Na^+^ and K^+^ concentrations relate. In order to understand how ion exchange between the extra- and intranuclear space takes place, we used isolated nuclei of HeLa cells. The isolated nuclei maintained their structural integrity (for at least one hour after isolation) according to FLIM data using benzothiazole orange (BO) (Fig. 4a), a fluorescent probe to nucleic acids [44,45]. The nuclei were incubated in media containing 10 mM Na^+^/100 mM K^+^ and 100 mM Na^+^/10 mM K^+^ that mimicked the cytoplasmic ion composition under normal and complete NKA inhibition conditions, respectively. In addition, to exclude possible active transport of the tested ions through the nuclear membrane [46], these experiments were also performed in the presence of 1 µM ouabain. In 10 mM Na^+^/100 mM K^+^ medium, the intranuclear Na^+^ and K^+^ ion concentrations were 17 ± 2 and 125 ± 12 mM, respectively, whereas in case of 100 mM Na^+^/10 mM K^+^ medium, the concentrations were 88 ± 20 and 11 ± 2 mM for Na^+^ and K^+^, respectively. In both cases, ouabain had no effect on the determined parameters (Fig. 4b). These data allow us to draw two conclusions. First, the values of calculated Na^+^ and K^+^ concentrations inside the nuclei in all cases are in equilibrium with the concentration of the tested ions in the extranuclear medium. Second, despite the possible presence of NKA in the nuclear membrane [46] ouabain did not affect the intranuclear concentrations of Na^+^ and K^+^.

**Fig. 4.**
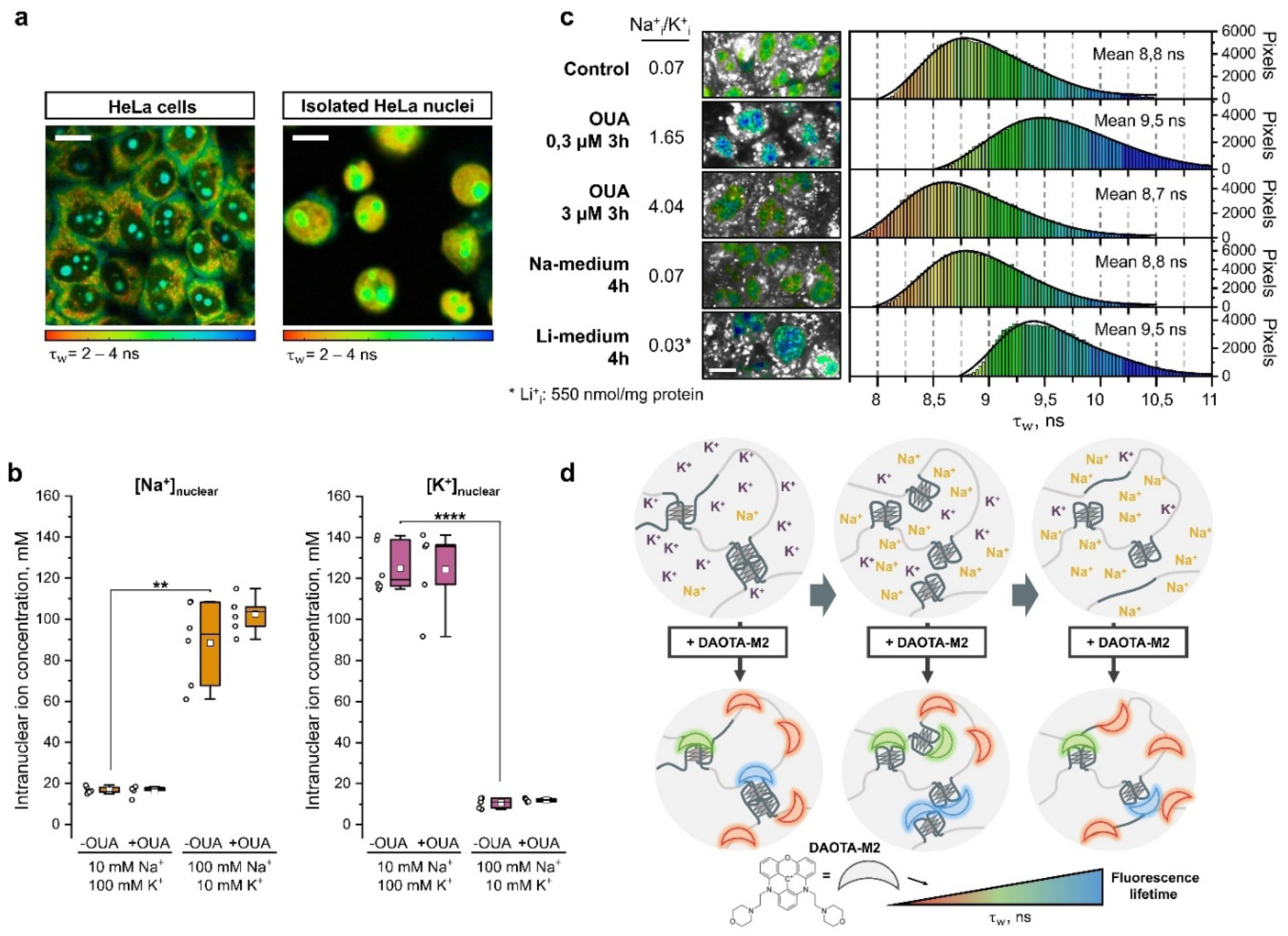
Monovalent metal cation imbalance in HeLa nuclei triggers G4 dynamics. **a.** Representative images of intact HeLa cells and isolated HeLa nuclei visualized using DNA-binding probe benzothiazole orange and FLIM. **b.** Concentration of Na^+^ and K^+^ in isolated HeLa nuclei upon changes in extranuclear concentration of these ions. The suspension of isolated nuclei shown in **a**, was incubated in 1 ml of 20 mM HEPES-imidazole, pH 7.8, 5 mM MgCl_2_, 0.5 mM DTT, 0.25 M sucrose, 1 mM EGTA, 15 mM ATP, containing different concentrations of Na^+^ and K^+^ in the presence or the absence of ouabain at 37 °C for 30 min. After incubation, the nuclei were washed three times with 1 ml of 0.1 M MgCl_2_ and then 1 ml of 5% TCA was added. The resulting suspension was centrifuged and the content of monovalent cations in the supernatant were determined. This content was normalized to the total volume of HeLa nuclei. Statistically significant differences were revealed using Welch-corrected ANOVA followed by post hoc Dunnet’s T3 test; \*\**p < 0.01*, \*\*\*\**p < 0.0001.* **с.** FLIM images of HeLa cells loaded with DAOTA-M2 recorded at 512×512 resolution (λ_ex_ =477nm, λ_em_=570–610 nm) and τ_w_ histograms from the nuclei area. Сells were incubated in DMEM supplemented with 0.1% FBS and 20 μM DAOTA-M2 for 24 h, after which they were treated with 0.3 or 3 μM ouabain (OUA) for 3 h or incubated in Na-/Li-medium for 4 h. Left side of the panel shows the intracellular Na^+^_i_/K^+^_i_ ratios that were registered under such conditions (for details see Fig. 1). Four different fields were analyzed to obtain histograms. Scale bars are 10 µm. **d.** Proposed mechanism of Na^+^_i_/K^+^_i_ ratio influence on G4s dynamics and their interaction with DAOTA-M2 in a living cell.

Based on this data, we assumed that intracellular concentrations of monovalent cations reflect their intranuclear concentrations and investigated the dynamics of DNA G4s in the nuclei of living cells in response to changes in intracellular monovalent cation content. For this purpose, we used FLIM of DAOTA-M2-loaded HeLa cells. Following P. Summers et al. [43] we used the intensity-weighted average lifetime τ_w_ of DAOTA-M2 as the most optimal G4 reporter. We increased intracellular Na^+^/K^+^ ratio by treating cells with 0.3 and 3 μM ouabain for 3 h. In addition, we loaded the cells with Li^+^ ions by incubating them in a medium in which all Na^+^ ions were replaced with Li^+^ ions. Incubation of HeLa cells in the presence of DAOTA-M2 (20 μM, 24 h) in the DMEM and Na-medium provided staining of their nuclei characterized by a normal τ_w_ distribution with mean value 8,8 ns (Fig. 4c). Subsequent incubation of HeLa cells in the presence of 0.3 μM ouabain (3 h) was accompanied by the increase in Na^+^_i_/K^+^_i_ ratio to 1.65 and in intranuclear τ_w_ to 9,5 ns, whereas in the presence of 3 μM ouabain (3 h) there were an increase in Na^+^_i_/K^+^_i_ ratio to 4.33 and a decrease in intranuclear τ_w_ to 8,7 ns compared to the control HeLa samples incubated in DMEM (Na^+^_i_/K^+^_i_ ratio 0.07; intranuclear τ_w_ 8,8 ns). HeLa cells incubated in Li-medium for 4 h were characterized by elevated intranuclear τ_w_ equal to 9,5 ns compared to control cells incubated in Na-medium (Na^+^_i_/K^+^_i_ ratio 0.07; intranuclear τ_w_ 8,9 ns) (Fig. 4 c). Notably, both controls (DMEM and Na-medium) provided the same intranuclear τ_w_ value. Thus, based on *in vitro* and *in cellulo* results, one can conclude that intracellular imbalance of monovalent cations affects the dynamics of DNA G4s by apparently changing not only their amount but also their topology (Fig. 4 d).

## Discussion

The idea that Na^+^ and K^+^ ions may participate in the regulation of gene expression in mammalian cells has been proposed earlier by some researchers [47]. The discovery of the phenomenon of Na^+^_i_/K^+^_i_-mediated Ca^2+^-independent regulation of gene expression naturally led to the assumption on the existence of intracellular molecular sensor(s) of monovalent cations [48]. We hypothesize that DNA G4s, whose structure formation depends on monovalent metal cations, could be at the same time such a sensor and a switch regulating gene expression. The regulatory role of Na^+^_i_/K^+^_i_ ratio in the formation and stabilization of DNA G4s was first pointed out by Sen and Gilbert, who termed this phenomenon “sodium-potassium switch” [49].

It is logical to assume that any stimulus affecting ion transport in the cell and leading to changes in the cytoplasmic ion composition should have its reflection on the ion composition of the nucleus. We have shown that the intranuclear ion composition changes proportionally depending on the concentration of monovalent cations in the extranuclear medium (Fig. 4b). Moreover, this effect was not dependent on ouabain that is inconsistent with previously published data according to which NKA is located in nuclear envelope and provide transport of Na^+^ and K^+^ between the cytoplasm and the intranuclear space [46]. Thus, we can consider that the nuclear ion composition reflects the ion composition of the cytoplasm.

According to the generally accepted conception, Li^+^ exerts a destabilizing effect on G4s structures [18]. Therefore, we suggest that the replacement of intracellular monovalent metal cations with Li^+^ could lead to alterations in the topology of G4s DNA and, consequently, to changes in genes transcription. Analyzing gene expression in response to an increase of the intracellular Na^+^_i_/K^+^_i_ ratio and Li^+^ loading of HeLa cells, we found that the expression patterns of *Fos*, *Jun*, *Zfp36*, and *Ptgs2* were similar under both stimuli. This suggests that such metal-dependent regulation is based on the same mechanism(s), which may be due to alterations in the structure and/or intermolecular interactions of G4s within the differentially expressed genes themselves or within other genes regulating them. Indeed, CD spectra of the two G4s from the *Fos* promoter that we tested (FOS_G4_1 and FOS_G4_3) showed a decrease in peak amplitude around 265 nm at Na^+^_i_/K^+^_i_ ratio > 1 and in the presence of 80 mM Li^+^, 40 mM K^+^, and 2 mM Na^+^ (Fig. 3b), which may indicate a transition from higher- to lower-ordered structures formed by G4s [50]. Other Na^+^_i_/K^+^_i_-dependent genes *Egr1*, *Atf3* (Fig. 2) were insensitive to the intracellular Li^+^ content; however, this does not exclude possible regulation of these genes by the alterations in G4s dynamics, because as was shown in our *in vitro* experiments different G4s have various sensitivity to monovalent metal cations (Fig. 3b). Thus, the structure type of telomeric G4 22AG changed in the presence of 80 mM Na^+^ and 20 mM K^+^, but not in the presence of 80 mM Li^+^, 40 mM K^+^, and 2 mM Na^+^ (Fig. 3b). Well-known examples of genes containing G4s in the promoter, such as *Kit* and *Myc*, changed their expression when exposed to Li^+^ (Fig.2c), but not when the intracellular Na^+^_i_/K^+^_i_ ratio was increased (Fig.2d). This is consistent with the CD spectroscopy data, according to which dramatic changes in the CD spectra of G4s from the promoters of these genes were observed in the presence of Li^+^ (Fig. 3b). Treatment of cells with G4-binding agent pyridostatin did not lead to a statistically significant change in the mRNA level of all the studied genes (Fig.2d), which is probably due to the different character of interaction of monovalent metal cations and chemical ligands with DNA G4s.

Since the *Fos* is an early response gene and encodes a pioneer transcription factor of the AP-1 family [12], it can be assumed that the Na^+^_i_/K^+^_i_-dependent mechanism of gene expression regulation is to some extent *Fos*-mediated in nature. Using the DNA-binding activity inhibitor of c-Fos/c-Jun T-5224, we showed that when binding to DNA c-Fos/c-Jun appears to suppress the expression of *Jun*, *Zfp36*, *Egr1*, *Atf3*, *Ptgs2*, *Kit* and also limits Na^+^_i_/K^+^_i_-mediated upregulation of *Zfp36* and *Egr1* (Fig. 1b). Apparently, the c-Fos/c-Jun binding to DNA provides negative feedback that limits Na^+^_i_/K^+^_i_-dependent induction of gene expression. Interestingly, the binding sites of AP-1 family transcription factors are enriched in G4-folded sequences [51], which may indicate a probable regulation of G4s dynamics near these sites by AP-1 binding. At the same time, no effect of T-5224 on *Fos* transcription was observed, even in ouabain-treated cells, which was unexpected, given the presence of AP-1 binding element in the promoter of this gene [14]. Thus, it may be concluded that *Fos* is a key participant in the Na^+^_i_/K^+^_i_-dependent regulation of gene expression in mammalian cells. We deduce that the “sensitivity” of this gene to monovalent metal cations may be determined by the presence of at least three G4s in its promoter structure, which have different topologies that can alter depending on the ion environment (Fig. 3a, b). This can play an important role in the development of various pathologies associated with a violation of the Na^+^_i_/K^+^_i_ ratio. In particular, in Alzheimer’s disease, a disruption of the Na^+^_i_/K^+^_i_ gradient in brain tissue is observed [7], due to inhibition of NKA by *β*-amyloid peptide [52–56]. According to our data, this should lead to a change in *Fos* expression. Indeed, data on *Fos* overexpression in the hippocampus in Alzheimer’s disease have been appearing for a long time [57–59]. Recent data indicate that in Alzheimer’s disease *Fos* is an associated cellular senescence gene in the astrocyte, microglia, and pericyte/endothelial cell subgroups [60]. Thus, in this case, the proposed relationship between the Na^+^_i_/K^+^_i_ ratio and the level of *Fos* expression is observed.

Although there are many methods for studying DNA G4s *in cellulo*, the use of most of them involves sample preparation, during which the integrity of the plasma membrane and, as a consequence, ion homeostasis is disrupted [61]. In this regard, to assess the effect of monovalent metal cations perturbations on the dynamics of DNA G4s in living cells, we used a fluorescent probe DAOTA-M2, which permeates the cell, in conjunction with FLIM [43]. The dynamics of G4s implies changes in at least three parameters: the number of formed G4s, their topology, and the interaction of G4s with other molecules, including oligomerization of these structures. Given the fact that DAOTA-M2 stacks on top of DNA G4s, it remains unclear whether its fluorescence lifetime depends on the topology of the DNA G4s. Since the studied DNA G4s have different structures, we characterized the fluorescence lifetime of DAOTA-M2 upon binding to these DNA G4s at different ratios of monovalent metal cations concentrations. First of all, we noted that the weighted average fluorescence lifetime (τ_w_) of DAOTA-M2 is longer upon binding to the parallel type of DNA G4s (FOS_G4_1 and MYC_G4_), which indicates the dependence of this parameter on the DNA G4s topology. For most DNA G4s incubated with DAOTA-M2, the weighted average fluorescence lifetime (τ_w_) of DAOTA-M2 was significantly lower in the presence of 100 mM Na^+^or Li^+^ compared to 100 mM K^+^. These results may arise from the G4/double-stranded DNA balance shifted towards double-stranded DNA, since DAOTA-M2 has significantly shorter fluorescence lifetimes in combination with double-stranded DNA [43]. The fact that drastic changes in monovalent metal cation conditions may affect G4s structure and stability *in vitro* has been well covered [18]. However, this phenomenon has little physiological relevance, since Na^+^ or Li^+^-only conditions are not observed in the cytoplasm of animal cells even in pathological cases. That was a reason why we have studied G4s binding to DAOTA-M2 in the media that resemble the intracellular Na^+^/K^+^ ratio in cells experiencing partial (20 mM K^+^, 80 mM Na^+^), (50 mM K^+^, 50 mM Na^+^), and complete (20 mM K^+^, 80 mM Na^+^) NKA inhibition. We observed that τ_w_ (DAOTA-M2) values in complexes with most G4 sequences were barely affected by the Na^+^/K^+^ ratio in the incubation medium. One exception was FOS_G4_3: in this case τ_w_ (DAOTA-M2) increased by approximately 2 ns (when 100 mM K^+^ was changed to 20 mM Na^+^, 80 mM K^+^) and then decreased proportionally to the Na^+^/K^+^ ratio in the media.

Surprisingly, when the Na^+^_i_/K^+^_i_ ratio was changed, a similar nonlinear pattern in dynamics of DNA G4s in living cells was observed. Thus, with an increase in the Na^+^_i_/K^+^_i_ ratio from 0.07 to 1.6 (in the presence of 0.3 µM of ouabain), τ_w_ (DAOTA-M2) in the entire population of HeLa cells nuclei increased by almost 0.7 ns. With a further increase in the Na^+^_i_/K^+^_i_ ratio to values of the order of 4.3 (in the presence of 3 µM of ouabain), τ_w_ (DAOTA-M2) was 0.2 ns less than in the control. Similar patterns observed *in vitro* (in the case of FOS_G4_3) and *in cellulo* can be explained, for example, by the competition of G4s formation processes and a decrease in their stability. According to recent concepts, DNA G4s also exist in the form of oligomers in the chromatin structure [62]. In this regard, with an increase in the Na^+^/K^+^ ratio, monomeric intramolecular G4s become prevailing over intermolecular G4s and G4s associations. Such processes should probably lead to an increase in the proportion of monomeric G4s *in vitro* or in the chromatin of a living cell, which should be accompanied by an augmentation of the DAOTA-M2 fluorescence lifetime due to an increase in the number of binding sites for the dye. On the other hand, at higher Na^+^/K^+^ ratios, a complete destabilization of G4s in favor of non-quadruplex forms, such as double-stranded or hairpin DNA, leads to a decrease in the number of G4s. Thus, these processes may compensate for and even outweigh the effect of G4s deoligomerization, which leads to a decrease in the DAOTA-M2 fluorescence lifetime. At the same time, when interpreting the data on the decrease of τ_w_ (DAOTA-M2) in the presence of 3 µM ouabain, it is worth considering the cytotoxic effect of ouabain at micromolar concentrations [63], which can initiate cell death processes accompanied by chromatin remodeling [64].

As an alternative G4s-destabilizing treatment, we loaded HeLa cells with Li^+^, hypothesizing that Li^+^ mimics an increased Na^+^_i_/K^+^_i_ ratio. Indeed, in both cases we observed an augmentation in mRNA levels of such Na^+^_i_/K^+^_i_-sensitive genes as *Fos*, *Jun*, *Zfp36*, and *Ptgs2*. However, only Li^+^ loading but not an elevation of the Na^+^_i_/K^+^_i_ ratio decreased *Kit* and *Myc* mRNA levels (Fig. 2c, d). Earlier, we observed similar effects in HUVECs [65]. Moreover, in a medium simulating the intracellular ion composition upon loading cells with Li^+^, we detected a dramatic change in the CD spectra for KIT_G4_ and MYC_G4_ (Fig. 3b). It should be also noted that both the accumulation of Li^+^ and the increase in the Na^+^_i_/K^+^_i_ ratio under the action of 0.3 µM ouabain in HeLa were accompanied by an identical increase in τ_w_ (DAOTA-M2) by 700 ps (Fig. 4c). Thus, our data support the assumption that DNA G4s can orchestrate the expression of certain genes upon alterations of the monovalent metal cation balance in animal cells.

Considering the involvement of DNA G4s in transcription regulation, their abundance in promoter regions, and sensitivity to monovalent cations, we assume that G4s are the “missing link” in the mechanism of gene regulation by monovalent metal cations. Apparently, metal cations affect the oligomeric state of DNA G4s, their topology, quantity, and the ability to interact with ligands, thus regulating the transcriptional activity of gene promoters.

## Acknowledgments

All measurements of the fluorescence lifetimes of dyes were conducted using the equipment of the Lomonosov Moscow State University Center for Picosecond Fluorescence Spectroscopy and Microscopy, supported by the Russian Science Foundation grant No. 25-75-00013. The analysis of parameter distributions using differential evolution methods was performed within the framework of the Russian Science Foundation (RSF) project No. 25-21-00375.

## Abbreviations

^1^H NMR: proton nuclear magnetic resonance
AP-1: activator protein 1
CRE: cAMP response element
CREB: cAMP response element-binding protein
DMEM: Dulbecco’s Modified Eagle Medium
FBS: fetal bovine serum
FLIM: fluorescence lifetime imaging microscopy
G4: G-qudruplex
HUVEC: human umbilical vein endothelial cells
MS: mass spectrometry
NKA: Na,K-ATPase
OUA: ouabain
PDS: pyridostatin
PQS: potential G-quadruplex-forming sequence
RCE: retinoblastoma control element
RVSMC: rat vascular smooth muscle cells
SIE: sis-inducible element
SRE: serum response element
SRF: serum response factor
TCF: ternary complex factor
TRE: 12-O-tetradecanoylphorbol-13-acetate response element

## Notes

### Competing Interest Statement

The authors have declared no competing interest.

## References

[1] N.J. Gerkau, R. Lerchundi, J.S.E. Nelson, M. Lantermann, J. Meyer, J. Hirrlinger, C.R. Rose, Relation between activity-induced intracellular sodium transients and ATP dynamics in mouse hippocampal neurons, J Physiol 597 (2019) 5687–5705. 10.1113/JP278658.

[2] K. Danilov, S. Sidorenko, K. Milovanova, E. Klimanova, L.V. Kapilevich, S.N. Orlov, Electrical pulse stimulation decreases electrochemical Na+ and K+ gradients in C2C12 myotubes, Biochemical and Biophysical Research Communications 493 (2017) 875–878. 10.1016/j.bbrc.2017.09.133.

[3] G. Sjøgaard, R.P. Adams, B. Saltin, Water and ion shifts in skeletal muscle of humans with intense dynamic knee extension, Am J Physiol 248 (1985) R190–196. 10.1152/ajpregu.1985.248.2.R190.

[4] I.L. Cameron, N.K.R. Smith, T.B. Pool, R.L. Sparks, Intracellular Concentration of Sodium and Other Elements as Related to Mitogenesis and Oncogenesis in Vivo, Cancer Res 40 (1980) 1493–1500.

[5] O.D. Lopina, D.A. Fedorov, S.V. Sidorenko, O.V. Bukach, E.A. Klimanova, Sodium Ions as Regulators of Transcription in Mammalian Cells, Biochemistry Moscow 87 (2022) 789– 799. 10.1134/S0006297922080107.

[6] A. Shatrova, E. Burova, N. Pugovkina, A. Domnina, N. Nikolsky, I. Marakhova, Monovalent ions and stress-induced senescence in human mesenchymal endometrial stem/stromal cells, Sci Rep 12 (2022) 11194. 10.1038/s41598-022-15490-2.

[7] V.M. Vitvitsky, S.K. Garg, R.F. Keep, R.L. Albin, R. Banerjee, Na+ and K+ ion imbalances in Alzheimer’s disease, Biochim Biophys Acta 1822 (2012) 1671–1681. 10.1016/j.bbadis.2012.07.004.

[8] S. V. Koltsova, Y. Trushina, M. Haloui, O.A. Akimova, J. Tremblay, P. Hamet, S.N. Orlov, Ubiquitous [Na+]i/[K+]i-Sensitive Transcriptome in Mammalian Cells: Evidence for Ca2+i-Independent Excitation-Transcription Coupling, PLoS ONE 7 (2012) e38032. 10.1371/journal.pone.0038032.

[9] L. Smolyaninova, S. Koltsova, S. Sidorenko, S. Orlov, Augmented gene expression triggered by Na+,K+-ATPase inhibition: Role of Ca2+i-mediated and −independent excitation-transcription coupling, Cell Calcium 68 (2017) 5–13. 10.1016/j.ceca.2017.10.002.

[10] M. Kappelmann, A. Bosserhoff, S. Kuphal, AP-1/c-Jun transcription factors: Regulation and function in malignant melanoma, European Journal of Cell Biology 93 (2014) 76–81. 10.1016/j.ejcb.2013.10.003.

[11] S. Taurin, N.O. Dulin, D. Pchejetski, R. Grygorczyk, J. Tremblay, P. Hamet, S.N. Orlov, c-Fos Expression in Ouabain-Treated Vascular Smooth Muscle Cells from rat Aorta: Evidence for an Intracellular-Sodium-Mediated, Calcium-Independent Mechanism, The Journal of Physiology 543 (2002) 835–847. 10.1113/jphysiol.2002.023259.

[12] S.C. Biddie, S. John, P.J. Sabo, R.E. Thurman, T.A. Johnson, R.L. Schiltz, T.B. Miranda, M.-H. Sung, S. Trump, S.L. Lightman, C. Vinson, J.A. Stamatoyannopoulos, G.L. Hager, Transcription Factor AP1 Potentiates Chromatin Accessibility and Glucocorticoid Receptor Binding, Molecular Cell 43 (2011) 145–155. 10.1016/j.molcel.2011.06.016.

[13] S. Bahrami, F. Drabløs, Gene regulation in the immediate-early response process, Advances in Biological Regulation 62 (2016) 37–49. 10.1016/j.jbior.2016.05.001.

[14] E. Tulchinsky, Fos family members: regulation, structure and role in oncogenic transformation, Histol Histopathol 15 (2000) 921–928. 10.14670/HH-15.921.

[15] M. Haloui, S. Taurin, O.A. Akimova, D.-F. Guo, J. Tremblay, N.O. Dulin, P. Hamet, S.N. Orlov, [Na+]i-induced c-Fos expression is not mediated by activation of the 5′-promoter containing known transcriptional elements, The FEBS Journal 274 (2007) 3557–3567. 10.1111/j.1742-4658.2007.05885.x.

[16] Y. Nakagawa, V. Rivera, A.C. Larner, A role for the Na/K-ATPase in the control of human c-fos and c-jun transcription., Journal of Biological Chemistry 267 (1992) 8785–8788. 10.1016/S0021-9258(19)50347-7.

[17] H.J. Lipps, D. Rhodes, G-quadruplex structures: in vivo evidence and function, Trends in Cell Biology 19 (2009) 414–422. 10.1016/j.tcb.2009.05.002.

[18] D. Bhattacharyya, G. Mirihana Arachchilage, S. Basu, Metal Cations in G-Quadruplex Folding and Stability, Frontiers in Chemistry 4 (2016). https://www.frontiersin.org/articles/10.3389/fchem.2016.00038 (accessed April 22, 2023).

[19] Y. Luo, M.L. Živković, J. Wang, J. Ryneš, S. Foldynová-Trantírková, L. Trantírek, D. Verga, J.-L. Mergny, A sodium/potassium switch for G4-prone G/C-rich sequences, Nucleic Acids Research 52 (2024) 448–461. 10.1093/nar/gkad1073.

[20] E.A. Venczel, D. Sen, Parallel and antiparallel G-DNA structures from a complex telomeric sequence, Biochemistry 32 (1993) 6220–6228. 10.1021/bi00075a015.

[21] G. Marsico, V.S. Chambers, A.B. Sahakyan, P. McCauley, J.M. Boutell, M.D. Antonio, S. Balasubramanian, Whole genome experimental maps of DNA G-quadruplexes in multiple species, Nucleic Acids Res 47 (2019) 3862–3874. 10.1093/nar/gkz179.

[22] R. Hänsel-Hertsch, M. Di Antonio, S. Balasubramanian, DNA G-quadruplexes in the human genome: detection, functions and therapeutic potential, Nat Rev Mol Cell Biol 18 (2017) 279–284. 10.1038/nrm.2017.3.

[23] J.L. Huppert, S. Balasubramanian, G-quadruplexes in promoters throughout the human genome, Nucleic Acids Res 35 (2007) 406–413. 10.1093/nar/gkl1057.

[24] Z.-H. Zhang, S.H. Qian, D. Wei, Z.-X. Chen, In vivo dynamics and regulation of DNA G-quadruplex structures in mammals, Cell & Bioscience 13 (2023) 117. 10.1186/s13578-023-01074-8.

[25] O.H. Lowry, N.J. Rosebrough, L. Farr, R. Randall J., Protein measurement with the Folin phenol reagent, Journal of Biological Chemistry 193 (1951) 265–275. 10.1007/978-94-007-0753-5_100521.

[26] S.N. Rampersad, Multiple Applications of Alamar Blue as an Indicator of Metabolic Function and Cellular Health in Cell Viability Bioassays, Sensors 12 (2012) 12347–12360. 10.3390/s120912347.

[27] C. Guilluy, L.D. Osborne, L. Van Landeghem, L. Sharek, R. Superfine, R. Garcia-Mata, K. Burridge, Isolated nuclei adapt to force and reveal a mechanotransduction pathway within the nucleus, Nat Cell Biol 16 (2014) 376–381. 10.1038/ncb2927.

[28] K.J. Livak, T.D. Schmittgen, Analysis of Relative Gene Expression Data Using Real-Time Quantitative PCR and the 2−ΔΔCT Method, Methods 25 (2001) 402–408. 10.1006/meth.2001.1262.

[29] O. Doluca, G4Catchall: A G-quadruplex prediction approach considering atypical features, Journal of Theoretical Biology 463 (2019) 92–98. 10.1016/j.jtbi.2018.12.007.

[30] Q. Zhai, C. Gao, J. Ding, Y. Zhang, B. Islam, W. Lan, H. Hou, H. Deng, J. Li, Z. Hu, H.I. Mohamed, S. Xu, C. Cao, S.M. Haider, D. Wei, Selective recognition of c-MYC Pu22 G-quadruplex by a fluorescent probe, Nucleic Acids Res 47 (2019) 2190–2204. 10.1093/nar/gkz059.

[31] S.A. Lizunova, V.B. Tsvetkov, D.A. Skvortsov, P.N. Kamzeeva, O.M. Ivanova, L.A. Vasilyeva, A.A. Chistov, E.S. Belyaev, A.A. Khrulev, T.S. Vedekhina, A.N. Bogomazova, M.A. Lagarkova, A.M. Varizhuk, A.V. Aralov, Anticancer activity of G4-targeting phenoxazine derivatives in vitro, Biochimie 201 (2022) 43–54. 10.1016/j.biochi.2022.07.001.

[32] A. Shivalingam, M.A. Izquierdo, A.L. Marois, A. Vyšniauskas, K. Suhling, M.K. Kuimova, R. Vilar, The interactions between a small molecule and G-quadruplexes are visualized by fluorescence lifetime imaging microscopy, Nat Commun 6 (2015) 8178. 10.1038/ncomms9178.

[33] A.N. Semenov, E.G. Maksimov, A.M. Moysenovich, M.A. Yakovleva, G.V. Tsoraev, A.A. Ramonova, E.A. Shirshin, N.N. Sluchanko, T.B. Feldman, A.B. Rubin, M.P. Kirpichnikov, M.A. Ostrovsky, A.N. Semenov, E.G. Maksimov, A.M. Moysenovich, M.A. Yakovleva, G.V. Tsoraev, A.A. Ramonova, E.A. Shirshin, N.N. Sluchanko, T.B. Feldman, A.B. Rubin, M.P. Kirpichnikov, M.A. Ostrovsky, Protein-Mediated Carotenoid Delivery Suppresses the Photoinducible Oxidation of Lipofuscin in Retinal Pigment Epithelial Cells, Antioxidants 12 (2023). 10.3390/antiox12020413.

[34] D.A. Fedorov, S.V. Sidorenko, A.I. Yusipovich, O.V. Bukach, A.M. Gorbunov, O.D. Lopina, E.A. Klimanova, Increased Extracellular Sodium Concentration as a Factor Regulating Gene Expression in Endothelium, Biochemistry Moscow 87 (2022) 489–499. 10.1134/S0006297922060013.

[35] E.A. Klimanova, S.V. Sidorenko, P.A. Abramicheva, A.M. Tverskoi, S.N. Orlov, O.D. Lopina, Transcriptomic Changes in Endothelial Cells Triggered by Na,K-ATPase Inhibition: A Search for Upstream Na+i/K+i Sensitive Genes, Int J Mol Sci 21 (2020) 7992. 10.3390/ijms21217992.

[36] Y. Aikawa, K. Morimoto, T. Yamamoto, H. Chaki, A. Hashiramoto, H. Narita, S. Hirono, S. Shiozawa, Treatment of arthritis with a selective inhibitor of c-Fos/activator protein-1, Nat Biotechnol 26 (2008) 817–823. 10.1038/nbt1412.

[37] A.M. Gorbunov, D.A. Fedorov, O.E. Kvitko, O.D. Lopina, E.A. Klimanova, FOS Promoter is Overactive Outside of Genome Context and Weakly Regulated by Changes in the Na+i/K+i Ratio, Biochemistry Moscow 90 (2025) 582–589. 10.1134/S0006297925600371.

[38] V.E. Yurinskaya, A.V. Moshkov, T.S. Goryachaya, A.A. Vereninov, Li/Na exchange and Li active transport in human lymphoid cells U937 cultured in lithium media, Cell Tiss. Biol. 8 (2014) 80–90. 10.1134/S1990519X1401012X.

[39] R. Rodriguez, S. Müller, J.A. Yeoman, C. Trentesaux, J.-F. Riou, S. Balasubramanian, A Novel Small Molecule That Alters Shelterin Integrity and Triggers a DNA-Damage Response at Telomeres, J. Am. Chem. Soc. 130 (2008) 15758–15759. 10.1021/ja805615w.

[40] K. Lu, H.-C. Wang, Y.-C. Tu, C.-C. Chang, P.-J. Lou, T.-C. Chang, J.-J. Lin, Suppressing c-FOS expression by G-quadruplex ligands inhibits osimertinib-resistant non-small cell lung cancer, J Natl Cancer Inst 115 (2023) 1383–1391. 10.1093/jnci/djad142.

[41] A. Kotar, B. Wang, A. Shivalingam, J. Gonzalez-Garcia, R. Vilar, J. Plavec, NMR Structure of a Triangulenium-Based Long-Lived Fluorescence Probe Bound to a G-Quadruplex, Angewandte Chemie International Edition 55 (n.d.) 12111–12542. 10.1002/anie.201606877.

[42] A. Shivalingam, A. Vyšniauskas, T. Albrecht, A.J.P. White, M.K. Kuimova, R. Vilar, Trianguleniums as Optical Probes for G-Quadruplexes: A Photophysical, Electrochemical, and Computational Study, Chemistry – A European Journal 22 (2016) 4129–4139. 10.1002/chem.201504099.

[43] P.A. Summers, B.W. Lewis, J. Gonzalez-Garcia, R.M. Porreca, A.H.M. Lim, P. Cadinu, N. Martin-Pintado, D.J. Mann, J.B. Edel, J.B. Vannier, M.K. Kuimova, R. Vilar, Visualising G-quadruplex DNA dynamics in live cells by fluorescence lifetime imaging microscopy, Nat Commun 12 (2021) 162. 10.1038/s41467-020-20414-7.

[44] T.L. Netzel, K. Nafisi, M. Zhao, J.R. Lenhard, I. Johnson, Base-Content Dependence of Emission Enhancements, Quantum Yields, and Lifetimes for Cyanine Dyes Bound to Double-Strand DNA: Photophysical Properties of Monomeric and Bichromomphoric DNA Stains, J. Phys. Chem. 99 (1995) 17936–17947. 10.1021/j100051a019.

[45] T. Biver, A. Boggioni, F. Secco, E. Turriani, M. Venturini, S. Yarmoluk, Influence of cyanine dye structure on self-aggregation and interaction with nucleic acids: A kinetic approach to TO and BO binding, Archives of Biochemistry and Biophysics 465 (2007) 90–100. 10.1016/j.abb.2007.04.034.

[46] C. Galva, P. Artigas, C. Gatto, Nuclear Na+/K+-ATPase plays an active role in nucleoplasmic Ca2+ homeostasis, J Cell Sci 125 (2012) 6137–6147. 10.1242/jcs.114959.

[47] S.N. Orlov, P. Hamet, Intracellular Monovalent Ions as Second Messengers, J Membrane Biol 210 (2006) 161–172. 10.1007/s00232-006-0857-9.

[48] E.A. Klimanova, S.V. Sidorenko, A.M. Tverskoi, A.A. Shiyan, L.V. Smolyaninova, L.V. Kapilevich, S.V. Gusakova, G.V. Maksimov, O.D. Lopina, S.N. Orlov, Search for Intracellular Sensors Involved in the Functioning of Monovalent Cations as Secondary Messengers, Biochemistry Moscow 84 (2019) 1280–1295. 10.1134/S0006297919110063.

[49] D. Sen, W. Gilbert, A sodium-potassium switch in the formation of four-stranded G4-DNA, Nature 344 (1990) 410–414. 10.1038/344410a0.

[50] P. Tóthová, P. Krafčíková, V. Víglaský, Formation of Highly Ordered Multimers in G-Quadruplexes, Biochemistry 53 (2014) 7013–7027. 10.1021/bi500773c.

[51] S. Lago, M. Nadai, F.M. Cernilogar, M. Kazerani, H. Domíniguez Moreno, G. Schotta, S.N. Richter, Promoter G-quadruplexes and transcription factors cooperate to shape the cell type-specific transcriptome, Nat Commun 12 (2021) 3885. 10.1038/s41467-021-24198-2.

[52] I.Y. Petrushanko, V.A. Mitkevich, A.A. Anashkina, A.A. Adzhubei, K.M. Burnysheva, V.A. Lakunina, Y.V. Kamanina, E.A. Dergousova, O.D. Lopina, O.O. Ogunshola, A.Y. Bogdanova, A.A. Makarov, Direct interaction of beta-amyloid with Na,K-ATPase as a putative regulator of the enzyme function, Sci Rep 6 (2016) 27738. 10.1038/srep27738.

[53] C.A. Dickey, M.N. Gordon, D.M. Wilcock, D.L. Herber, M.J. Freeman, D. Morgan, Dysregulation of Na+/K+ ATPase by amyloid in APP+PS1 transgenic mice, BMC Neuroscience 6 (2005) 7. 10.1186/1471-2202-6-7.

[54] C. Kairane, R. Mahlapuu, K. Ehrlich, M. Zilmer, U. Soomets, The effects of different antioxidants on the activity of cerebrocortical MnSOD and Na,K-ATPase from post mortem Alzheimer’s disease and age-matched normal brains, Curr Alzheimer Res 11 (2014) 79–85. 10.2174/15672050113106660179.

[55] L.-N. Zhang, Y.-J. Sun, S. Pan, J.-X. Li, Y.-E. Qu, Y. Li, Y.-L. Wang, Z.-B. Gao, Na^+^-K^+^- ATPase, a potent neuroprotective modulator against Alzheimer disease, Fundam Clin Pharmacol 27 (2013) 96–103. 10.1111/fcp.12000.

[56] E.P. Barykin, I.Y. Petrushanko, S.A. Kozin, G.B. Telegin, A.S. Chernov, O.D. Lopina, S.P. Radko, V.A. Mitkevich, A.A. Makarov, Phosphorylation of the Amyloid-Beta Peptide Inhibits Zinc-Dependent Aggregation, Prevents Na,K-ATPase Inhibition, and Reduces Cerebral Plaque Deposition, Front Mol Neurosci 11 (2018) 302. 10.3389/fnmol.2018.00302.

[57] D.L. Marcus, J.A. Strafaci, D.C. Miller, S. Masia, C.G. Thomas, J. Rosman, S. Hussain, M.L. Freedman, Quantitative neuronal c-fos and c-jun expression in Alzheimer’s disease, Neurobiol Aging 19 (1998) 393–400. 10.1016/s0197-4580(98)00077-3.

[58] A.J. Anderson, B.J. Cummings, C.W. Cotman, Increased Immunoreactivity for Jun- and Fos-Related Proteins in Alzheimer’s Disease: Association with Pathology, Experimental Neurology 125 (1994) 286–295. 10.1006/exnr.1994.1031.

[59] W. Lu, R. Mi, H. Tang, S. Liu, M. Fan, L. Wang, Over-expression of c-fos mRNA in the hippocampal neurons in Alzheimer’s disease, Chin Med J (Engl) 111 (1998) 35–37.

[60] D. Sha, J. Zhang, X. Fang, X. Wang, X. He, X. Shu, Expression and potential regulatory mechanism of cellular senescence-related genes in Alzheimer’s disease based on single-cell and bulk RNA datasets, Front. Neurosci. 19 (2025). 10.3389/fnins.2025.1595847.

[61] M. Di Antonio, R. Rodriguez, S. Balasubramanian, Experimental approaches to identify cellular G-quadruplex structures and functions, Methods 57 (2012) 84–92. 10.1016/j.ymeth.2012.01.008.

[62] I. Frasson, V. Pirota, S.N. Richter, F. Doria, Multimeric G-quadruplexes: A review on their biological roles and targeting, International Journal of Biological Macromolecules 204 (2022) 89–102. 10.1016/j.ijbiomac.2022.01.197.

[63] O.D. Lopina, A.M. Tverskoi, E.A. Klimanova, S.V. Sidorenko, S.N. Orlov, Ouabain-Induced Cell Death and Survival. Role of α1-Na,K-ATPase-Mediated Signaling and [Na+]i/[K+]i-Dependent Gene Expression, Front. Physiol. 11 (2020). 10.3389/fphys.2020.01060.

[64] W. Park, S. Wei, B.-S. Kim, B. Kim, S.-J. Bae, Y.C. Chae, D. Ryu, K.-T. Ha, Diversity and complexity of cell death: a historical review, Exp Mol Med 55 (2023) 1573–1594. 10.1038/s12276-023-01078-x.

[65] O.E. Kvitko, D.A. Fedorov, S.V. Sidorenko, O.D. Lopina, E.A. Klimanova, Accumulation of Li+ Ions Triggers Changes in FOS, JUN, EGR1, and MYC Transcription in the LiCl-Treated Human Umbilical Vein Endothelial Cells (HUVEC), Biochemistry Moscow 89 (2024) 1844–1850. 10.1134/S0006297924100146.

