## Supplementary information for "Alteration in the intracellular Na^+^/K^+^ ratio regulates gene expression through the shift of G-quadruplex dynamics in living cells"

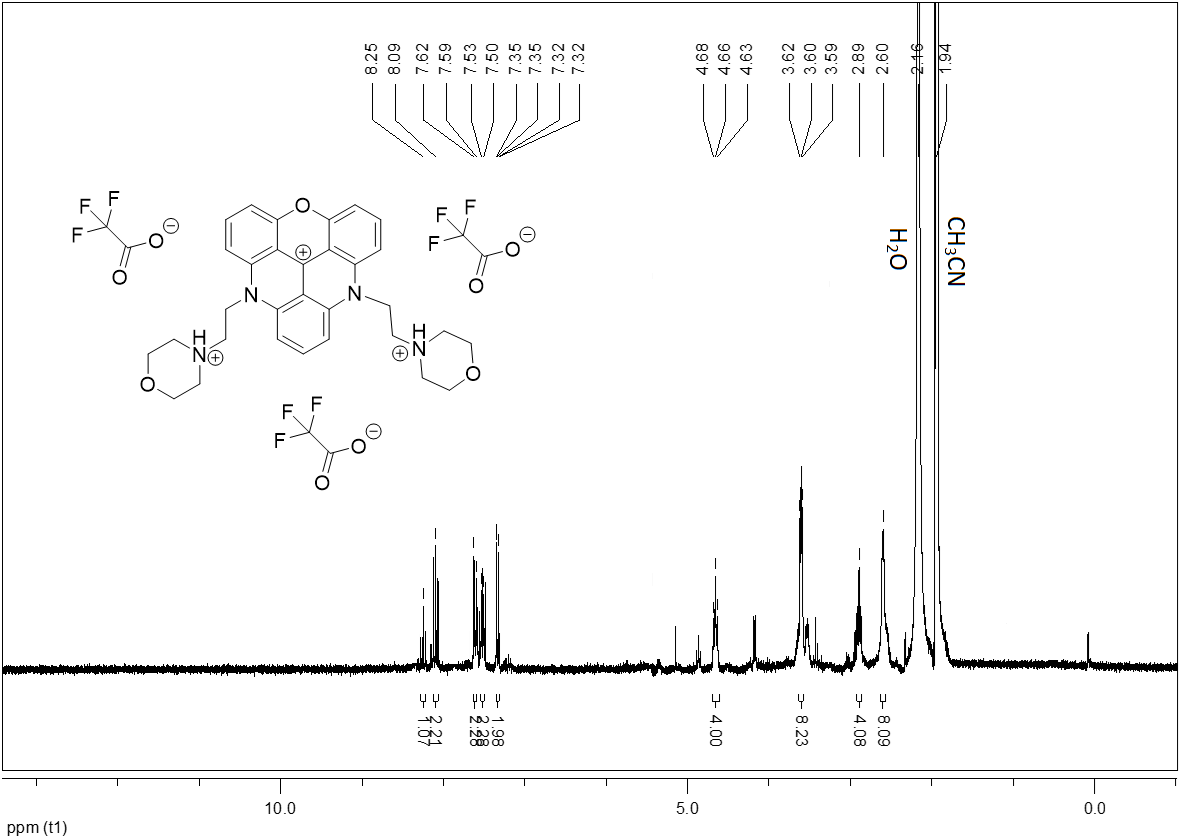

Fig. S1. ^1^H NMR spectrum of DAOTA-M_2_ was recorded on a Bruker Fourier 300 spectrometer (USA). ^1^H NMR (300 MHz, CD_3_CN): δ 8.25 (t, *J* = 8.7 Hz, 1H), 8.09 (dd, *J* = 8.8 Hz, *J* = 8.2 Hz, 2H), 7.61 (d, *J* = 8.9 Hz, 2H), 7.51 (d, *J* = 8.6 Hz, 2H), 7.33 (dd, J = 8.2 Hz, J = 0.5 Hz, 2H), 4.66 (t, *J* = 7.0 Hz, 4H), 3.63-3.56 (m, 8H), 2.89 (t, *J* = 7.0 Hz, 4H), 2.63-2.56 (m, 8H).

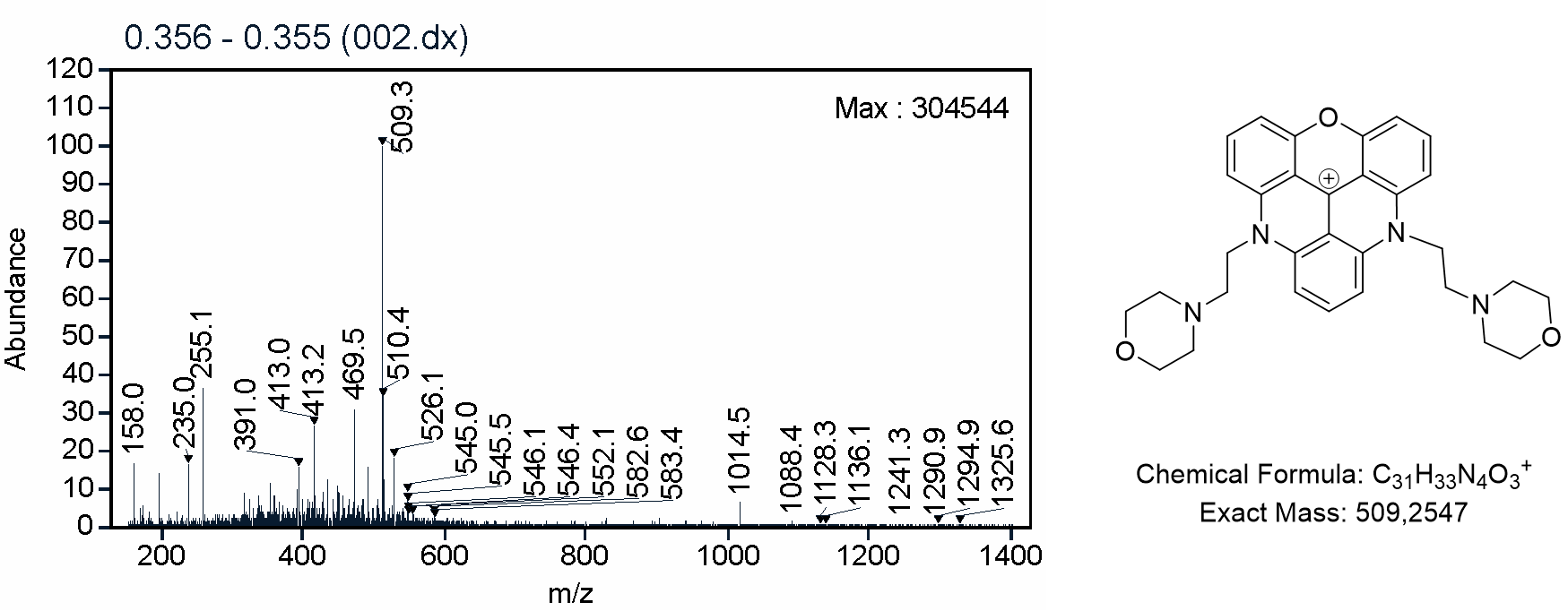

Fig. S2. MS spectrum of DAOTA-M_2_ was recorded on Agilent 6340 G2447A Ion Trap LC/MS System (USA) in continuous flow direct sample infusion (positive ion mode).

Table S1. Intracellular Na^+^ and K^+^ content in HeLa cells after 24 h incubation in control (DMEM) or in the presence of 10 µM pyridostatin.

|  | Na^+^_i_, nmol/mg protein | K^+^_i_, nmol/mg protein |
| --- | --- | --- |
| control | 73 ± 4 | 1015 ± 63 |
| 10 $\mu$M pyridostatin | 77 ± 8 | 971 ± 18 |

Data from six biological replicates (n=6) are represented as mean ± SD.
